# Structure and organization of full-length Epidermal Growth Factor Receptor in extracellular vesicles by cryo-electron tomography

**DOI:** 10.1101/2024.11.25.625301

**Authors:** Monica Gonzalez-Magaldi, Anuradha Gullapalli, Ophelia Papoulas, Chang Liu, Adelaide Y.-H. Leung, Luqiang Guo, Axel Brilot, Edward M. Marcotte, Zunlong Ke, Daniel J. Leahy

## Abstract

We report here transport of the Epidermal Growth Factor Receptor (EGFR), Insulin Receptor, 7-pass transmembrane receptor Smoothened, and 13-pass Sodium-iodide symporter to extracellular vesicles (EVs) for structural and functional studies. Mass spectrometry confirmed the transported proteins as the most abundant in EV membranes, and the presence of many receptor-interacting proteins demonstrates the utility of EVs for characterizing membrane protein interactomes. Cryo-electron tomography of EGFR-containing EVs reveals that EGFR forms clusters in the presence of EGF with a ∼3 nm gap between the inner membrane and cytoplasmic density. EGFR extracellular regions do not form regular arrays, suggesting that clustering is mediated by the intracellular region. Subtomogram averaging of the EGFR extracellular region (ECR) yielded a 15 Å map into which the crystal structure of the ligand-bound EGFR ECR dimer fits well. These findings refine our understanding of EGFR activation, clustering, and signaling, and they establish EVs as a versatile platform for structural and functional characterization of human membrane proteins in a native-like environment.

**Significance Statement:** Atomic or near-atomic resolution structural studies of proteins embedded in cell membranes have proven challenging. We show that transporting integral membrane proteins to cell-derived extracellular vesicles enables structural and functional studies of human membrane proteins in a native membrane environment. We have used this approach to visualize an active form of full-length Epidermal Growth Factor Receptor (EGFR) and show that it forms clusters in the membrane and projects its cytoplasmic signaling domains ∼3 nm away from the membrane surface. EGFR is essential for normal development, but abnormal EGFR activity is associated with several human cancers and is the target of many anticancer therapies. Our studies refine current models of how ligand binding to EGFR transmits signals across cell membranes.

## Introduction

Structural studies of soluble fragments or detergent-solubilized forms of integral membrane proteins have yielded enormous insight into diverse biological processes (1–3). For example, crystal structures of the extracellular region (ECR) of the Epidermal Growth Factor Receptor (EGFR), a Receptor Tyrosine Kinase (RTK) composed of a ligand binding ECR, a single transmembrane region, and an intracellular kinase domain, revealed a large domain rearrangement in the ECR when ligand is bound (4–7). This rearrangement exposes buried surfaces that mediate formation of canonical dimers of the EGFR ECR. This ECR dimer in turn promotes formation of an asymmetric dimer of intracellular kinase regions that is essential for signaling (8, 9).

These observations are consistent with the long-held model that EGFR is activated by ligand-dependent dimerization (10) but fail to explain all aspects of EGFR behavior. For example, EGFR has been shown to form multimers in the absence of ligand (11–18) and higher-order oligomers in the presence of ligand (19–21). In addition, distinct EGFR ligands stimulate different cellular responses in otherwise similar conditions(22–24). This phenomenon, known as biased agonism, has been attributed both to different ligands modulating EGFR ECR dimer structure in a way that is communicated to the intracellular kinase (25–27) and to different ligand binding and EGFR dimerization kinetics observed for different ligands (28). These explanations are not necessarily mutually exclusive.

Understanding whether or how the EGFR ECR communicates structural information to the ICR beyond dimerization and the nature and importance of non-canonical EGFR oligomers would be greatly aided by visualization of intact forms of EGFR in native cell membranes. Several recent studies have used single-particle cryo-electron microscopy to examine full-length or near full-length forms of EGFR or EGFR homologs solubilized in detergents, nanodiscs, or peptidiscs (27, 29, 30). These studies have revealed ordered ECR dimers that are largely consistent with crystal structures, but considerable flexibility between ECRs and intracellular regions (ICRs) precluded resolution of ICR structures.

To enable structural and functional studies of EGFR and other integral membrane proteins in cell-derived membranes we developed a system to transport human membrane proteins to Virus-like Particles (VLPs). VLPs are non-infectious enveloped viral particles ∼60-150 nm in diameter that bud off from cell membranes and whose membranes are enriched in viral co-expressed viral proteins (31, 32). Recent cryo-electron tomography (cryo-ET) and single-particle cryo-electron microscopic (cryo-EM) studies of proteins on viral surfaces have resulted in subnanometer and near-atomic resolution structures, respectively, demonstrating the potential for structural studies of proteins expressed in VLP membranes (33).

To transport human membrane proteins to VLPs, we fused epitope-binding modules to the regions of viral surface proteins that mediate transport to viral membranes and showed that these fusion proteins (i) transport to VLPs and (ii) “pull” epitope-tagged human membrane proteins including EGFR, Insulin Receptor (InsR), Smoothened (Smo), and the Sodium-iodide symporter (NIS) into VLPs. Strategic placement of rhinovirus 3C protease recognition sites allows removal of both the fusion protein extracellular regions and the peptide epitopes. Hoffman, Bjorkman, and colleagues recently reported that appending a 110 amino-acid sequence containing binding regions for ESCRT and ALIX proteins to the SARS-CoV-2 spike protein resulted in formation of spike-containing virus-like vesicles they term enveloped VLPs (eVLPs) (34). We find that appending this ESCRT and ALIX binding region (EABR) to EGFR or other human single- and multi-pass integral membrane proteins results in production of vesicles containing the EABR-labeled protein. As these vesicles contain no viral proteins, we will refer to them as extracellular vesicles (EVs).

Using EGFR as an example, we show that EVs are a simplified and advantageous system for structural and functional studies of human membrane proteins in native bilayers. Mass spectrometric analysis of EGFR-containing EVs shows enrichment in many known EGFR-interacting proteins, demonstrating that EVs can be used to characterize membrane protein interactomes, analogous to the Virotrap approach for cytoplasmic proteins (35). Cryo-electron Tomography (cryo-ET) analyses of EGFR-containing EVs in the presence of excess EGF reveal that EGFR forms clusters in EV membranes but that the EGFR ECRs do not form regular contacts, suggesting that clustering is mediated by the EGFR intracellular region (ICR). A ∼3 nm gap is also observed between the inner membrane surface and cytoplasmic density indicating that EGFR kinase domains are projected away from the membrane in the active state.

Subtomogram averages of the 150 kDa EGFR ECR dimer resulted in 15 Å density maps into which the crystal structure of dimer of the EGFR-ECR:EGF complex fits well.

## Results

### Transport of human membrane proteins to VLPs

To create molecules that bind epitope-tagged proteins of interest and transport them to virus-like particles (VLPs), Strep-tactin (SA) and single-chain Fv (scFv) fragments of antibodies that recognize Hemagglutin (HA) (36) or FLAG (37) peptides were substituted for the extracellular region of Influenza Neuraminidase (Figure 1A). To allow for simple detection of these molecules, which we term “transporters,” and increase accessibility of the scFv regions to epitopes on target proteins, glycine-serine (Gly-Ser) linkers and tandemly repeated myc tags were inserted between Neuraminidase and scFV sequences, and a triple repeat of FLAG sequences was inserted between Neuraminidase and SA. A rhinovirus 3C protease recognition site was also added to the juxtamembrane region of each transporter to enable proteolytic removal of extracellular regions after VLP formation (Figure 1A).

**Figure 1.**
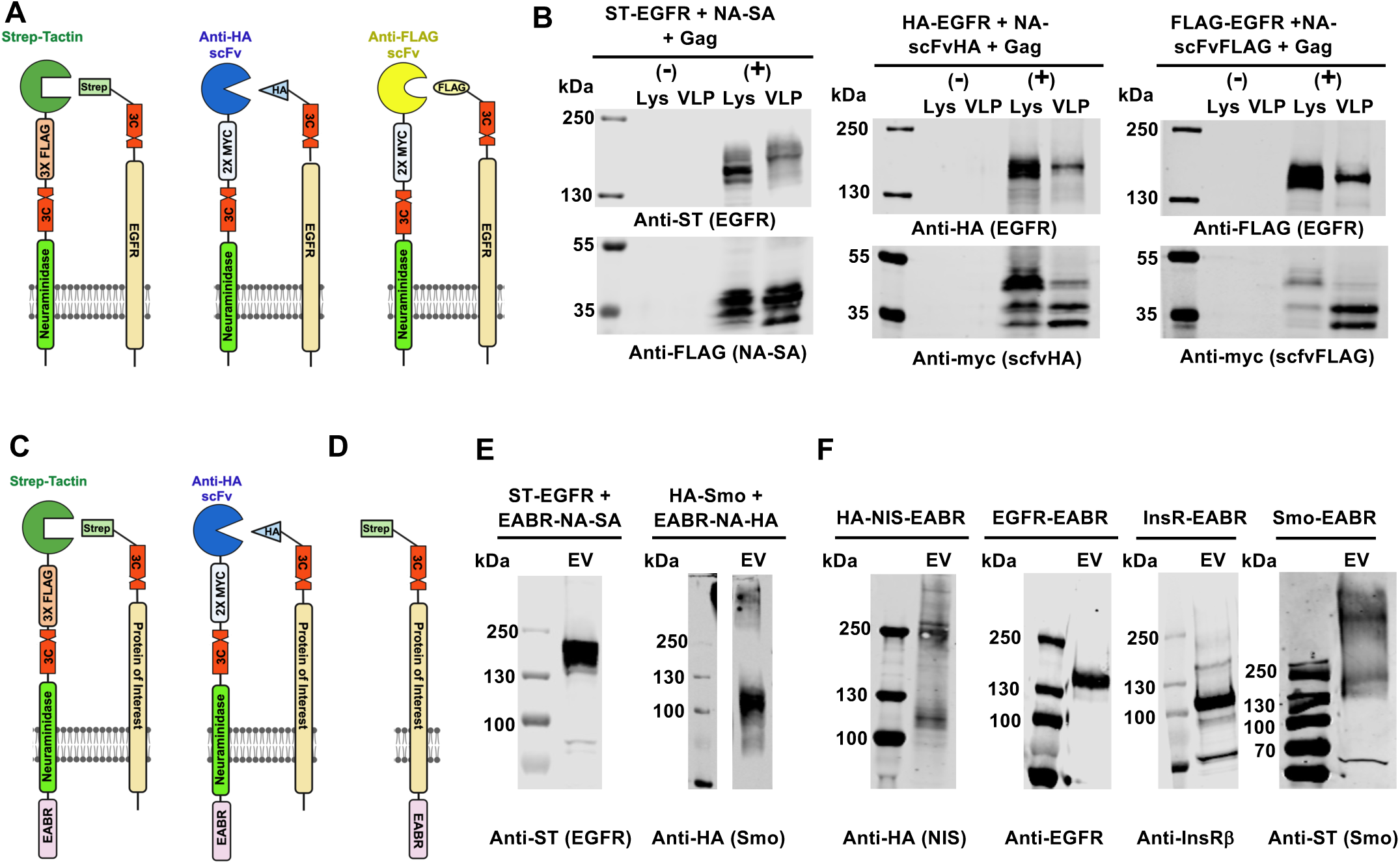
Transport of human proteins to VLPs and EVs. (A) Schematic diagram of proteins containing the N-terminal region of influenza neuraminidase fused to peptide binding module including Strep-Tactin (SA), a single-chain Fv targeting a hemagglutinin peptide (Anti-HA scFv), or a single-chain Fv targeting the FLAG peptide (Anti-FLAG scFv). FLAG and myc epitopes were included in these proteins to enable detection, and a Rhinovirus 3C protease recognition site (3C) included to enable proteolytic removal of transport protein extracellular regions. Labels read in the N to C direction. (B) Western blots of HEK 293 cell either not transfected (-) or transfected (+) with the MLV Gag gene, a transporter protein, and an EGFR gene with sequences encoding an HA, FLAG, or Strep-tag II (ST) epitope at its N-terminus. Lanes loaded with cell lysates (Lys) and with VLPs following purification by ultracentrifugation through a sucrose cushion (VLP). (C) Schematic diagram of proteins containing the ESCRT and Alix binding region (EABR) fused to the N-terminal region of influenza neuraminidase followed by a peptide binding module. (D) Schematic diagram of a protein of interest with the EABR fused to its C-terminus to promote transport to EVs and a Strep-tag II epitope fused to its N-terminus to enable purification of EVs using a Strep-Tactin affinity column. (E) Western blot analysis of sucrose cushion purified EVs derived from Expi293 cells transfected with the transporters diagrammed in (D) along with either Strep-tag II labeled EGFR (left) or HA- labeled Smoothened (Smo). (F) Western blot analysis of sucrose cushion purified EVs derived from Expi293 cells transfected with genes encoding the Sodium-iodide symporter (NIS) (left), EGFR (center-left), Insulin Receptor (InsR) (center-right), or Smoothend (Smo) (right) with an EABR tag fused to their C-termini. The probing antibody is indicated below each blot.

Each of the HA, FLAG, and SA transporters expressed well in CHO and HEK293 cells and led to VLP formation from HEK 293T cells when co-transfected with a gene encoding the Murine leukemia virus Gag protein (Figure 1B) (38). Initial purification and concentration of VLPs was achieved by centrifuging VLP-containing media through a 20% sucrose cushion and resuspension of the VLP-containing pellet. When co-expressed with a gene encoding human EGFR with a cognate epitope tag on its N-terminus, each transporter mediated transport of EGFR to VLPs (Figure 1B). Cryo-electron micrographs of EGFR-containing VLPs showed the presence of membrane-associated particles with size consistent with EGFR in the majority of VLPs, with empty VLPs presumably arising from cells expressing Gag but not EGFR proteins. Addition of Rhinovirus 3C protease to VLPs resulted in cleavage of both the transporter extracellular regions and the N-terminal epitope tags on EGFR (Figure S1), leaving EGFR behind as the most abundant membrane protein with sizeable extra- and intra- cellular regions. These transporters transported tagged but not untagged proteins to VLPs produced in HEK 293T cells. When untagged EGFR was expressed in Expi293 cells at ∼6-fold higher levels than HEK 293T cells, however, the transporter was not needed for EGFR to enter VLPs suggesting that transporters and epitope tags may not be needed if the target protein expresses at high levels, consistent with prior results producing vesicles from cells overexpressing proteins (39, 40) (Figure S2).

### EABR tags target human membrane proteins to eVLPs

Hoffman, Bjorkman, and colleagues recently demonstrated that addition of a 110 amino-acid segment containing ESCRT and Alix binding regions (EABR) to the C-terminus of the SARS-CoV-2 spike protein resulted in exocytosis of spike-containing vesicles (34). We appended their EABR sequence, which also contain an endocytosis prevention motif (41), to the cytoplasmic N-termini of the SA and HA transporter proteins, EGFR, Insulin Receptor (InsR), the 7-pass transmembrane receptor Smoothened (Smo), and the 13-pass transmembrane Sodium-iodide symporter (NIS) and observed transport of each molecule to EVs when transfected into Expi293 cells (Figure 1C-F). Furthermore, addition of EABR to the C-terminus of EGFR and InsR did not inhibit their activation as judged by anti-phosphotyrosine Western blots (Figure S3). In the event that the EABR tag affects the structure, function, or distribution of the target molecule, the EABR-based transporters can be used. The ability of the EABR-based transporter and EABR-labeled EGFR to produce and enter EVs (Figure 2) indicates that the EABR tag is effective in both orientations relative to the membrane and when separated from the transmembrane region by greater than 500 amino acids.

**Figure 2.**
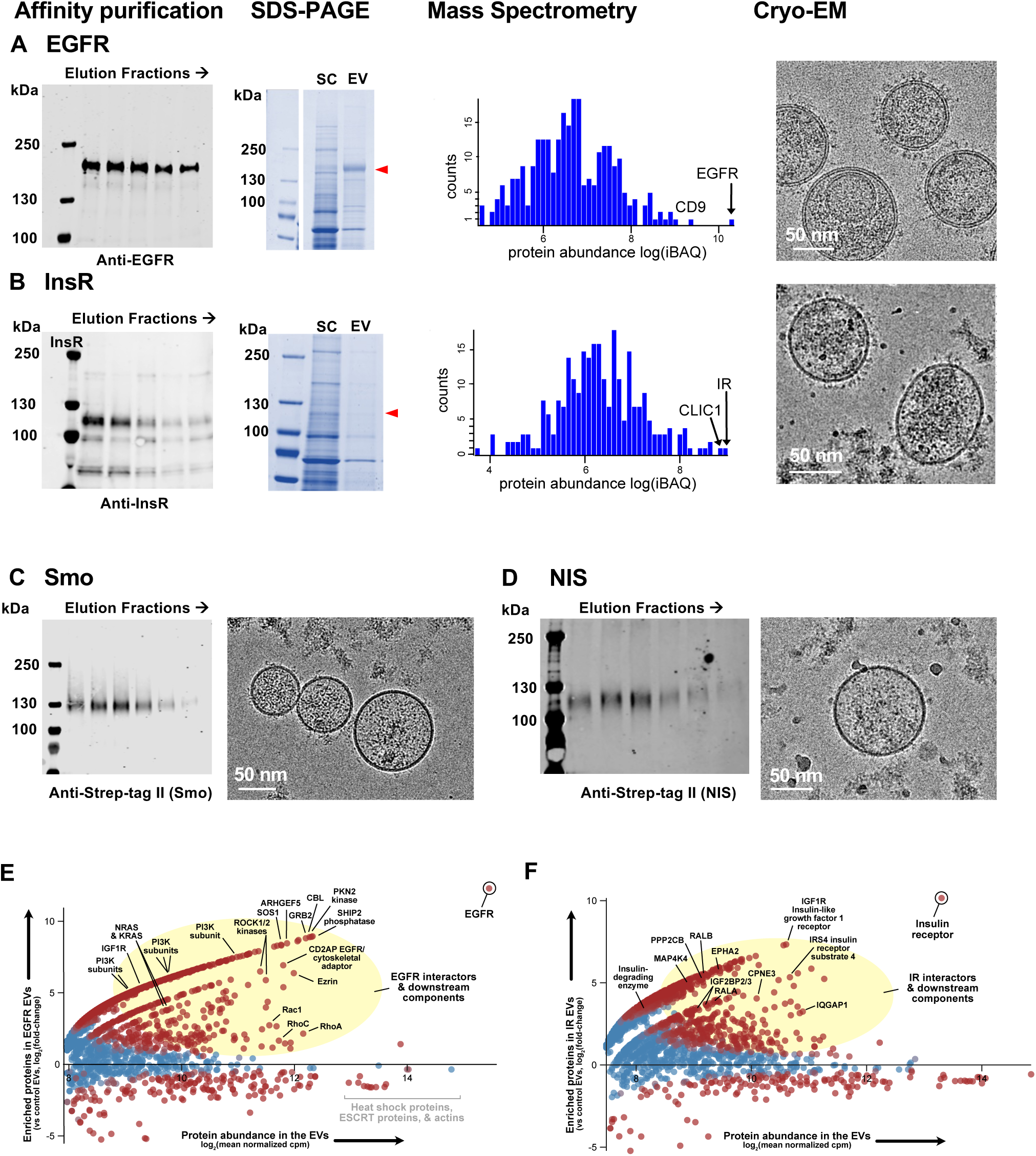
Affinity purification and characterization of EVs. Western blot of elution fractions from a Strep-Tactin column (left). Coomassie Blue stained SDS-PAGE analysis (center-left), mass spectrometric iBAQ analysis (center-right), and cryo-electron micrographs (right) of affinity-purified EVs containing Strep-tag II labeled (A) EGFR and (B) Insulin Receptor (IR). Elution fractions from a Strep-Tactin column (left) and cryo- electron micrographs (right) from Strep-tag II labeled (C) Smoothened (Smo) and (D) the Sodium-iodide Symporter (NIS). Probing antibodies are indicated below each blot. Plot of the log(Fold-change) (logFC) in the mass spectrometric counts of proteins in EVs produced by transfection with (E) EGFR-EABR or (F) Insulin Receptor-EABR relative to EVs without either receptor. The position of EGFR (E) and Insulin Receptor (F) are indicated along with the positions of several proteins known to interact with each receptor. A full list of abbreviations and interacting proteins can be found in Supplemental Information.

### Purification and characterization of EVs

Initial purification of VLPs and EVs from conditioned medium by centrifugation through a 20% sucrose cushion led to recovery of VLPs/EVs as judged by Western blot and cryo-electron microscopy (cryo-EM), but the micrographs indicated the presence of sufficient levels of contaminants to complicate electron-microscopic studies. A Strep-Tag II sequence (42) followed by a rhinovirus 3C protease recognition site (43) was thus added to the N-terminus of EABR-labeled EGFR, and EGFR-containing EVs further purified using a Strep-Tactin (SA) affinity resin (Figure 1C) (44). This affinity-purification step yielded EVs of sufficient purity for cryo-electron microscopic studies (Figure 2A) and was also used to purify InsR-, Smo-, and NIS-containing EVs (Figure 2B-D).

Intensity-based abundance quantification (iBAQ) (45) of mass spectrometric results obtained from EGFR-containing EVs following affinity purification indicate that the most abundant membrane protein is EGFR (Figure 2A). In addition, many known EGFR-interacting proteins were enriched in EGFR-containing EVs relative to non-EGFR-containing EVs (Figure 2E). Many of the enriched EGFR-interacting proteins, including Grb2, Shc1, PLCγ1, Cbl, and PI3K, bind phosphorylated EGFR suggesting that overexpression or clustering during EV production led to phosphorylation of EGFR (46).

Indeed, Western blot analysis of purified EGFR-containing EVs showed basal phosphorylation of EGFR that increased following addition of EGF (Figure S2). These results demonstrate that mass spectrometric analysis of EVs is an effective approach to characterizing the interactome of integral membrane proteins.

### Cryo-electron tomography of EGFR in EVs

EGF was added to EGFR-containing EVs prior to addition to electron microscope grids and plunge-freezing in liquid ethane.

Tomograms were calculated from tilt series collected on a Titan Krios equipped with a Gatan K3 detector and Biocontinuum energy filter. Tomograms revealed EVs with diameters of 88 ± 26 nm and surface particles of the size and shape expected for ligand-bound EGFR extracellular region dimers (Figures 3 and S4-S5). Visualization of whole EVs from tomograms can be challenging due to distortions arising from EVs abutting the edge of frozen ice and from the “missing wedge” of data, which leads to poorer resolution in the direction perpendicular to the plane of the EM grid. The program ISONET (47) was thus used to minimize missing wedge effects and reconstruct an isotropic view of EVs for a set of EVs entirely contained in vitreous ice (Figure 3A-C). EGFR dimers generally appear in clusters with a ∼3 nm gap between the intracellular membrane and intracellular density in both reconstructed and unreconstructed tomograms (Figure 3). Individual EGFRs are occasionally observed (Figures 3F-H and Figure S6) onto which the crystal structure of the EGFR:EGF complex dimer can be placed and with intracellular density consistent with the asymmetric EGFR kinase dimer projected ∼3 nm from the inner membrane surface.

**Figure 3.**
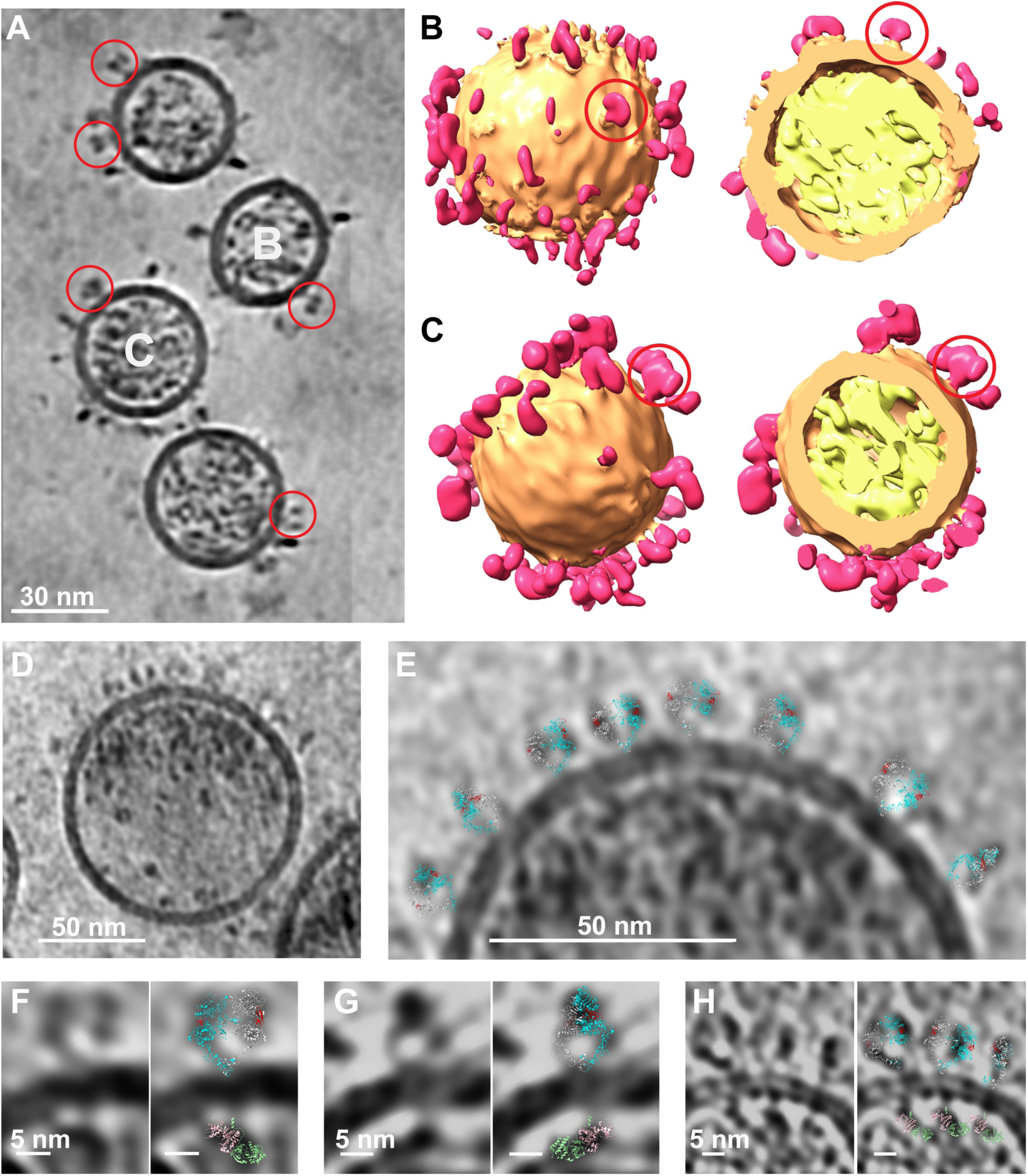
Tomograms of EGFR-containing EVs. (A) Section of tomogram of EGFR-containing EVs following application of ISONET to compensate for the missing wedge. Individual EGFR extracellular regions circled in red. (B-C) Enlarged views of two EVs from panel A showing the extracellular region (pink), membrane (tan), and intracellular region (yellow). Several EGFR extracellular regions are circled in red. (D) A section of a tomogram of an EGFR-containing EV is shown. (E) An enlarged region of the tomographic section shown in panel D is shown with ribbon diagrams of the EGFR extracellular region dimer (RCSB 3NJP; EGF is colored red and the EGFR subunits are colored cyan and light gray) manually positioned in the tomogram. (F-H) Images of individual EGFR molecules from tomograms of EGFR-containing EVs are shown alongside the same image with the crystal structures of the EGFR extracellular region dimer colored as in panel E and the asymmetric EGFR kinase dimer (RCSB 3GOP; one kinase subunit is colored light pink and the other light green).

To examine the arrangement of EGFRs in clusters, tomographic sections just outside EV surfaces in which cross-sections of EGFR clusters are present were inspected.

These sections show irregular arrangements of EGFR ECRs indicating that EGFR clustering is likely mediated by EGFR ICRs (Figure 4). Although isolated EGFR ICRs are occasionally observed, the ICRs of EGFRs in clusters are generally part of an amorphous assembly of density projected ∼3 nm below the inner membrane. To characterize this density, pie-shaped regions of a two-dimensional section of an EV tomogram were selected that either contained a cluster of EGFRs or were receptor-free. The average radial density of these regions was calculated and confirm accumulation of cytoplasmic density underneath EGFR clusters and the presence of a ∼3 nm gap between the inner membrane and EGFR cytoplasmic region (Figure 5A). Subtomogram averages of volumes from regions of 4 comparably-sized EVs that either contain a cluster of EGFRs or are receptor-free also demonstrate a concentration of cytoplasmic density at sites of EGFR clusters that is separated from the inner membrane by ∼3 nm (Figure 5B-D).

**Figure 4.**
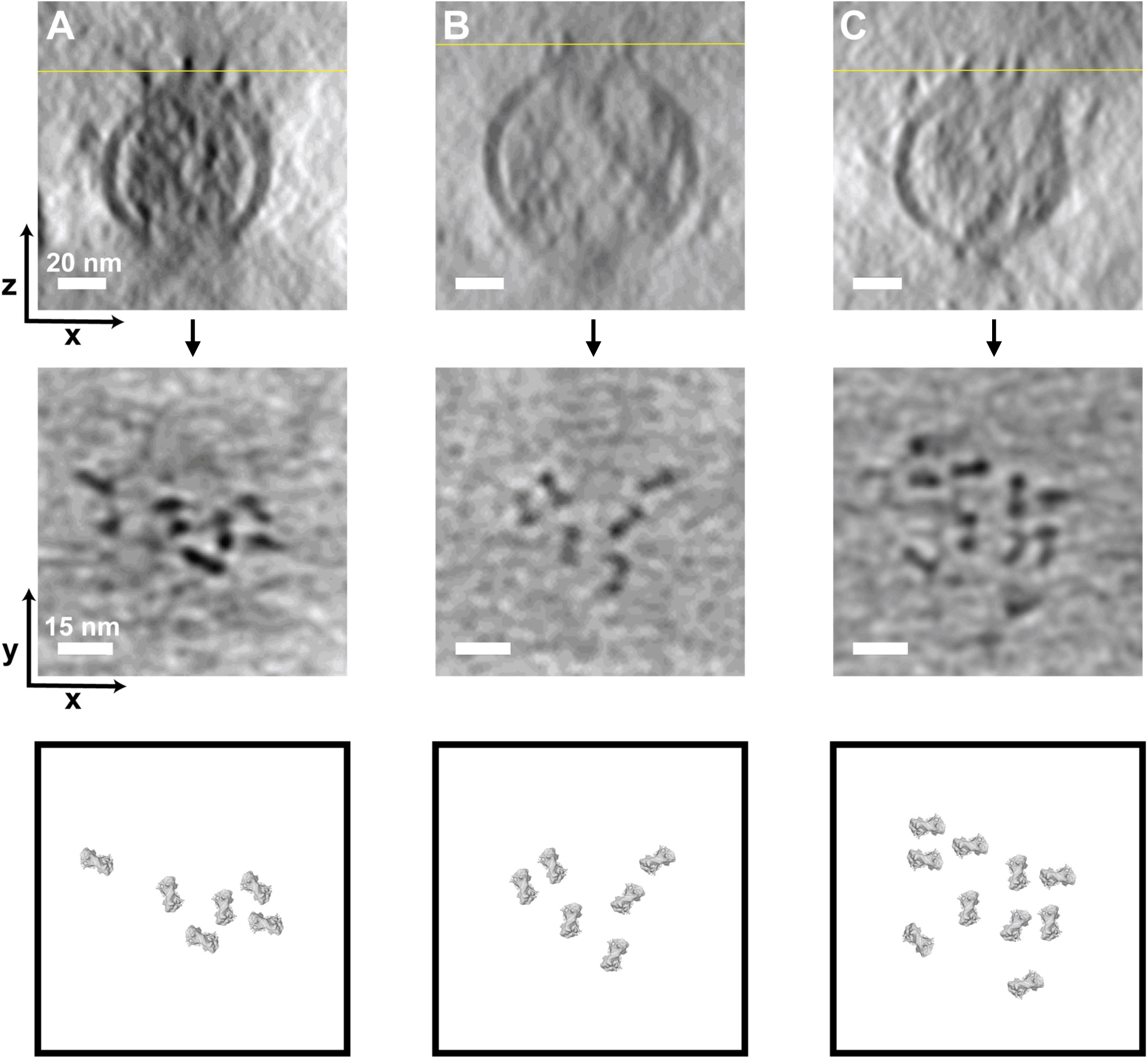
The extracellular regions of clustered EGFRs do not form regular arrays. Top panels: Tomographic sections of EVs fully embedded in ice, viewed perpendicular to the plane of the EM grid. Yellow lines indicate the approximate positions of the corresponding x-y planes shown in the middle panels. Middle panels: Tomographic sections of the same tomograms in the top row but viewed in the plane of the EM grid near the edges of the EVs. Bottom panels: Top-down views of cross-sections of the EGFR dimer derived from the crystal structure (RCSB: 3NJP), which are bowtie-shaped and ∼4.5 x 11 nm, positioned as in the middle panels.

**Figure 5.**
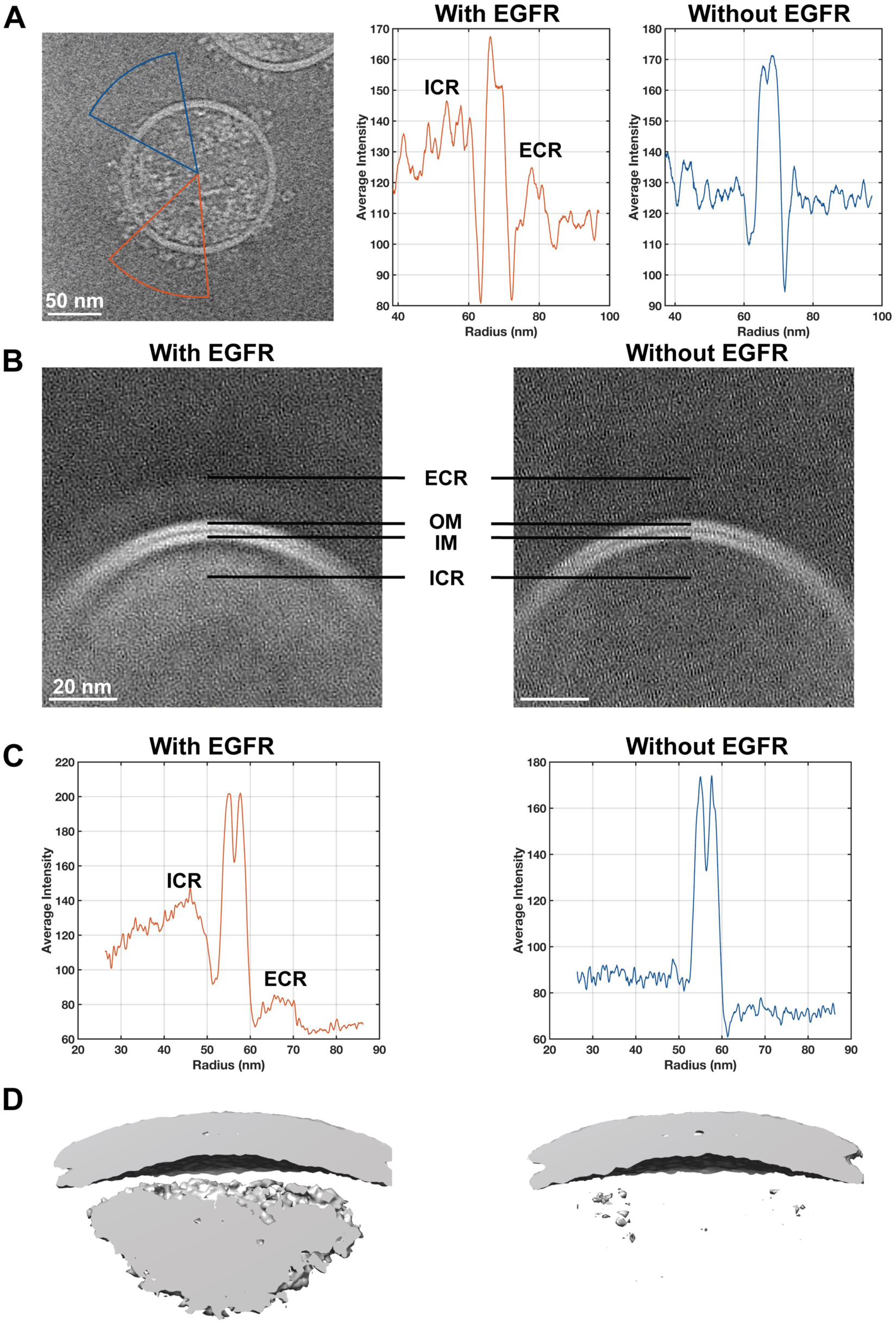
Characterization of intravesicular density. (A) 2D tomographic section (left) of an EGFR-containing EV showing regions with (orange) and without (blue) an EGFR cluster. The average radial density of these regions is shown in the middle and right panels. (B) 2D sections of a 3D subtomogram average of regions of 4 comparably-sized EVs that contain EGFR (left) or are devoid of EGFR (right). The extracellular region (ECR), outer membrane (OM), inner membrane (IM), and intracellular region (ICR) are indicated. Averaged radial density (C) and isosurface view (D) of the subtomogram averages shown in panel B.

The 31 amino-acid EGFR juxtamembrane region has been divided into an N-terminal 21 amino-acid segment (Juxtamembrane-A or JMA) and a C-terminal 10 amino-acid segment (Juxtamembrane-B or JMB) also called the “latch” because it forms an extended contact with another kinase in the active asymmetric EGFR kinase dimer (48, 49). If the JMA region adopts a random coil structure, it would likely not be rigid enough to project the EGFR kinase domains 3 nm away from the membrane. We thus believe the best explanation for this gap is that the JMA region forms a homodimeric parallel coiled-coil-like structure, which would be almost exactly 3 nm for a sequence of this length. Such a structure is predicted by AlphaFold 3 for a homodimer of the EGFR transmembrane and juxtamembrane regions (Figure S7) (50).

The appearance of recognizable EGFR ECRs in tomograms of EGFR-containing EVs led us to try structure determinations by both single-particle and subtomogram averaging. Initial attempts to determine the EGFR ECR structure by single-particle averaging of particles picked from cryo-EM projection images failed, however, which may stem from the overlap of multiple receptors in projection images of EGFR clusters. We thus focused our efforts on subtomogram averaging and were able to generate a 15 Å map of the EGFR ECR from 843 of 1602 manually picked particles (Figure 6A and Figure S8). These maps, on which C2 symmetry was imposed but otherwise used no template information from prior structures, faithfully reproduce features of the EGFR:EGF extracellular region crystal structure (RCSB 3NJP (51)) with the addition of bulges at sites of N-linked glycosylation (Figure 6A). Examination of subtomogram average classes from rejected particles fails to reveal classes in which the domain IV “legs” exhibit movements observed in a cryo-EM structure of EGFR in peptidiscs (27) and molecular dynamics simulations (52), which may indicate that the presence of the bilayer membrane stabilizes the EGFR ECR domain IV position observed in crystal structures (51).

**Figure 6.**
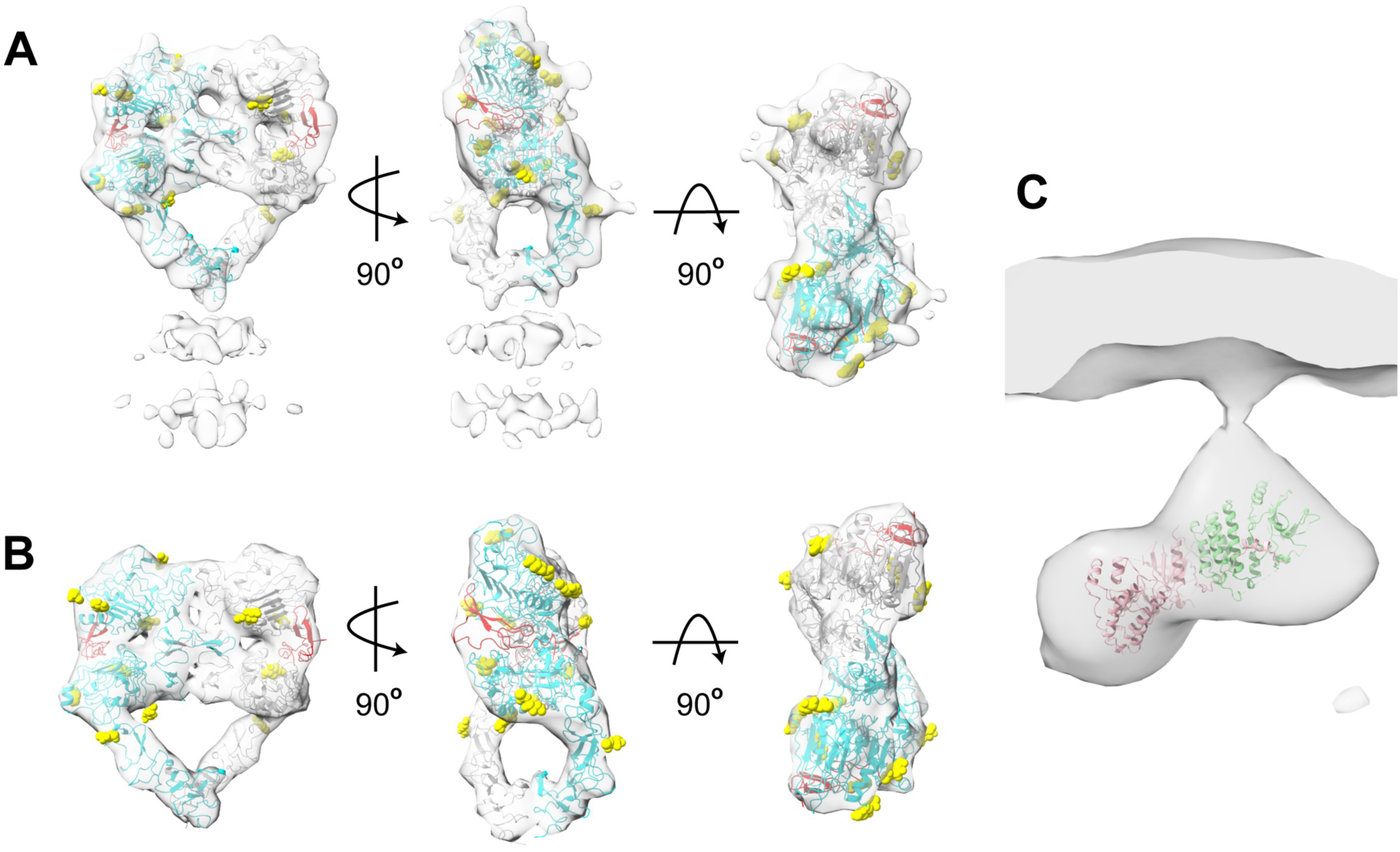
Subtomogram averages of the EGFR extracellular region in EVs strongly resemble the crystal structure. (A) Orthogonal views of the crystal structure of a dimer of the EGFR extracellular region complexed with EGF (RCSB 3NJP; the EGFR subunits are colored cyan and light gray and EGF is red) are shown fit to the 15 Å subtomogram average map of the EGFR extracellular regions on the surface of EVs. Sites of potential N-linked glycosylation are shown in yellow spheres of either N-acetyl glucosamines modeled in the crystal structure or Asparagine residues at consensus glycosylation sites. (B) Orthogonal views of the 15 Å map generated from the crystal structure of EGFR dimers bound to EGF as shown in panel A). (C) Subtomogram average of the intracellular regions of 16 isolated EGFRs with a ribbon the crystal structure of the EGFR asymmetric kinase dimer (RCSB 3GOP) shown with one light pink and one light green kinase subunit.

Owing to the scarcity of isolated EGFRs in EV membranes, the merging of intracellular density within EGFR clusters, and the likely association of EGFR interaction partners with EGFR ICRs, it proved difficult to identify well-defined and circumscribed EGFR ICR density. Sixteen such ICRs were identified, however, and when averaged produced a map consistent with the EGFR asymmetric kinase dimer (8) with a ∼3 nm stalk connecting it to the inner membrane surface (Figure 6B).

## Discussion

Previous structural studies of EGFR and related receptors have led to many valuable insights into EGFR function, but key questions remain. In particular, it is unclear whether and how structural information beyond dimerization is communicated from the extracellular to intracellular regions of the receptor as well as the nature and what roles EGFR multimers other than the active-state dimer might play in signaling. The ability to determine subnanometer or near-atomic resolution structures of viral envelope proteins by cryo-ET or cryo-EM, respectively, suggested that if EGFR or other human membrane proteins could be transported into VLP envelopes it would enable visualization of their structure and organization in native cell membranes (33, 53, 54).

We thus developed a “transporter” system for transporting epitope-tagged human membrane proteins to VLPs by attaching epitope binding modules to portions of viral surface proteins naturally transported to viral membranes. These transporters transported tagged but not untagged proteins to VLPs produced in HEK 293T cells, but when untagged EGFR was expressed in Expi293 cells at ∼6-fold higher levels the transporter was not needed for EGFR to enter VLPs suggesting that transporters and epitope tags may not be needed if the target protein expresses at high levels. In the midst of this work, Hoffmann, Bjorkman, and colleagues reported that appending a 110 amino-acid EABR sequence to the C-terminus of the SARS-CoV-2 spike led to efficient production of eVLPs containing the spike protein (34). We appended this EABR to EGFR, Insulin Receptor, the 7-pass transmembrane protein Smoothened, and the 13- pass transmembrane protein NIS and find efficient production of EABR-tagged protein containing EVs regardless of the EABR orientation or separation from the transmembrane region by up to 500 amino acids. The EABR system did not inhibit the activity of either EGFR or InsR as judged by phosphorylation assays and has the advantages that co-transfection with viral genes is not needed to produce vesicles and that virtually all EVs contain the desired protein. For these reasons the EABR system has become our method of choice although the transporter system has the advantages that (i) native or nearly native protein can be studied should the EABR tag affect function or distribution of the target protein and (ii) altering the ratio of target protein to transporter DNA in transfections may allow varying the concentration of the desired protein in EV/VLPs.

Mass spectrometric analysis of EGFR-containing EVs indicates that EGFR is greater than 10-fold more abundant than the next most abundant membrane protein as judged by iBAQ (45). We expected that the high abundance of EGFR in EV membranes would simplify particle identification in low-resolution cryo-electron tomograms, and a large majority of particles on the extravesicular surface in tomograms of EVs containing complexes of EGFR and EGF are indeed of the expected shape and size of EGFR-bound EGFR ECR dimers (51). These EGFR particles generally occur in clusters, but the EGFR ECRs do not form regular interactions, suggesting that cluster formation is mediated by EGFR ICRs, which in this case includes the EABR sequence at its C-terminus that may also promote clustering. The absence of EGFR ECR dimer-mediated multimers is almost certainly the case in physiological settings as, assuming physiological concentrations of EGFR are on the order of ∼100 receptors/µm^2^ (55)or ∼3 EGFRs per 100 nm diameter EV, EGFR concentrations in EVs are typically greater than 10-fold above physiological concentrations.

An unexpected feature of EGFR clusters is the presence of a ∼3 nm gap between the inner membrane and a layer of intracellular density that is occasionally spanned by weak tubes of density opposite EGFR ECRs. The JMA region of active EGFR dimers has been predicted to form an antiparallel coiled-coil based on NMR data (26), but such a structure could not explain the 3 nm gap we observe. We believe that the best explanation for the 21 amino-acid JMA region stably and rigidly projecting the EGFR kinase domains 3 nm away from the membrane is through formation a homodimeric parallel coiled-coil-like structure, and such a structure is predicted for the transmembrane and juxtamembrane regions by AlphaFold 3 (50). If a feature of inactive EGFR is separation of the transmembrane regions (55), an antiparallel coiled-coil of JMA regions may be favored in this state.

The observation of this gap raises the question of what its functional role might be. One possibility is suggested by the electrostatic model of EGFR activation in which interactions between a positively charged surface of the EGFR kinase and the negatively charged inner membrane surface inhibit EGFR activity in the absence of ligand and activation of the kinase requires release from the membrane (56). Formation of a coiled-coil of the JMA region in active EGFR dimer may thus be a mechanism to release inhibitory interactions between the EGFR kinase and the inner membrane surface. Support for this model comes from the observation that when the EGFR intracellular region was targeted to the inner membrane surface by a myristoylation signal it not highly active unless a GCN4 coiled-coil region was inserted between the myristoylation site and the kinase region with the objective of forcing dimerization and disrupting interaction between the kinase domain and membrane (26). Insertion of a GCN4 zipper in this case may have led to kinase activation by mimicking the structure and role of the JMA region in active EGFR.

Although cryo-electron microscopic studies of viral membrane proteins have reached subnanometer resolution for both tomographic and single particle studies, it was not clear that such resolutions could be obtained for proteins as small as the ∼150 kDa EGFR ECR dimer and its ∼72 kDa intracellular kinase dimer. Indeed, our attempts to produce single-particle averages of EGFR extracellular regions from projection images of EGFR-containing EVs have failed so far, presumably owing in part to overlap of adjacent receptors in projection images. Tomograms of EGFR-containing EVs contain recognizable EGFR extracellular regions that are separable, however, and subtomogram averages of 843 particles yields a density map with 15 Å resolution by the 0.143 FSC criterion. The crystal structure of the EGF-bound EGFR dimer fits into this map extremely well (Figure 6), indicating that crystal structures of isolated EGFR ECRs are an accurate reflection of the dominant conformation of EGFR ECRs in intact receptors in membrane bilayers.

The absence of additional density surrounding the EGFR ECR in subtomogram averages is consistent with EGFR ECRs not forming a conserved interdimer interaction.

Had a specific higher-order contact between EGFR ECR dimers been prevalent, density reflecting this contact would have appeared in our subtomogram averages. Bulges in density do occur, but they are invariably at sites of N-linked glycosylation. We also do not observe flexibility in the disposition of the domain IV relative to domain III, which was observed in single particle averages of EGFR dimers in peptidiscs (27) and molecular dynamics simulations of singly-ligated EGFR dimers (52). Although the C-terminal regions of domain IV come into close contact near the membrane, they appear to enter the membrane ∼30 Å apart, which is comparable to the distance between the ordered N-terminal residues in an NMR structure of an EGFR transmembrane region dimer (57) and may restrict tilting of EGFR ECRs in the direction parallel to the entry points.

In addition to facilitating structural studies of membrane proteins in host-cell membranes, EVs provide an excellent opportunity to characterize interaction partners of targeted membrane proteins by mass spectrometry. Most mass spectrometric approaches to identification of interaction partners rely on chemical crosslinking or persistence of interactions through detergent lysis and immunoprecipitation, each of which can lead to artifacts (58). Analysis of VLP or EVs rather than cells greatly reduces the number of background proteins and avoids many of these artifacts, and moreover, enrichment of putative interactors can be quantified in comparison to control EVs to provide a measure of confidence (Figure 2E-F). Our preliminary mass spectrometric analyses identified a large number of known EGFR interaction partners in EGFR-containing EVs, confirming that VLPs provide an excellent system to characterize the interactome of integral membrane proteins. A more detailed analysis of the EGFR interactome in active and inactive states will be reported elsewhere.

In this manuscript we report development of a VLP-transporter system and use of the EABR system of Hoffmann, Bjorkman, and colleagues (34) to stimulate production of extracellular vesicles containing human integral membrane proteins for structural and functional studies. These approaches differ from previous uses of EVs for structural studies in that vesicle formation is stimulated by engaging ESCRT proteins, the protein of interest is specifically targeted to vesicles, and use of sonication or chemical methods to produce vesicles is not needed (39, 40). We show that general features of integral membrane protein structure and organization can be inferred from inspection of individual proteins in tomograms and that proteins as small as the ∼150 kD EGFR ECR dimer can be imaged at near nanometer resolution using subtomogram averaging.

Mass spectrometric characterization of the proteomes of VLPs containing target proteins also provides a straightforward approach to identification of the interactome of integral membrane proteins. These results demonstrate that transport of human integral membrane proteins to EVs represents a promising approach to facilitate structural and function studies of human integral membrane proteins in native cell environments.

## Methods

### Construction of Transporters and tagged membrane proteins

pBS-CMV-gagpol was a gift from Patrick Salmon (Addgene plasmid #35614; http://n2t.net/addgene:35614; RRID:Addgene_35614). The polymerase gene, superfluous for VLP production, was deleted from pBS-CMV by PCR (59–61). To construct expression plasmids encoding Strep-tag II binding transporters, a synthetic gene-block encoding the N-terminal region of Influenza A virus Neuraminidase (residues 1-47 including transmembrane domain residues 7-35) (62) followed by a 5-residue Gly-Ser linker, an HRV 3C protease recognition site (43), a 5-residue Gly-Ser linker, three FLAG epitopes, a 5-residue Gly-Ser linker, and streptavidin (1-159 aa) with 3 mutations that confer higher affinity to Strep-tag II (Strep-tactin) (63), was inserted into the pBS-CMV backbone. To construct an HA epitope binding transporter, sequences encoding the N-terminal region of Influenza A virus Neuraminidase (residues 1-47 including transmembrane domain residues 7-35) followed by 5-residue Gly-Ser linker, an HRV 3C protease recognition site, a 15-residue Gly-Ser linker, two myc epitopes, an 8-residue Gly-Ser linker, and the anti-HA scFv from pCMV-15F11-HA-mEGFP, which was a gift from Tim Stasevich (Addgene plasmid #129590 ; http://n2t.net/addgene:129590 ; RRID:Addgene_129590) (36), was inserted into the pBS-CMV backbone. The FLAG epitope binding transporter was contructed similarly to the HA transporter except that anti-FLAG heavy and light chain sequences derived a plasmid encoding anti-FLAG M2 antibody (37), which was a gift from Joost Snijder (Addgene plasmid #175359; #175358 http://n2t.net/addgene:175359; RRID:Addgene_175359 or 8) were used to create the scFv region. Expression plasmids directing expression of full-length Insulin Receptor (InsR) and EGFR with and without N-terminal HA and FLAG epitopes are described elsewhere (64), pcDNA HA-NIS was a gift from Nancy Carrasco. Sequences encoding a Strep-tag II epitope followed by a Gly-Ser linker and a HRV 3C site at the N-termini of EGFR, InsR, NIS and SMO were added to each respective expression plasmids, and versions of these clones with sequences encoding the EABR sequence at their C-termini were constructed using the sequence reported by Hoffmann, Bjorkman, and colleagues (34). To construct a negative control for analysis of interactomes by mass spectrometry, the coding sequence of EGFR transmembrane region (TM) was cloned into pUE1-TSP-BFLF1, a gift from Britt Glaunsinger (Addgene plasmid #162657 ; http://n2t.net/addgene:162657 ; RRID:Addgene_162657) replacing the BFLF1 sequence by the TM and adding to the N-terminus tandemly repeated Strep-tag II epitopes followed by a 5x Gly-Ser linker, an Alfa tag, a 5x Gly-Ser linker, an HRV 3C region and a 5x Gly-Ser linker. The 31 aa transmembrane region is followed by the EABR coding sequence.

*Transfection of CHO, HEK293T and Expi293 cells.* CHO-K1 cells (ATCC CCL-61) and HEK293T (ATCC CRL-3216) cells were maintained in DMEM/F12 medium supplemented with 5-10% inactivated FBS. To test protein expression, cells were seeded in 6 well plates for transfection with PEI (Polyethyleneimine ‘Max’ linear, MW 25 000 1 mg/ml) using a 3:1 w/w PEI:DNA ration for each plasmid to be tested. Cells were washed with PBS 24h post transfection and lysed in RIPA buffer (50 mM Tris pH 8.0, 150 mM NaCl, 1% v/v NP40, 0.5% w/v sodium deoxycholate, 0.1% w/v SDS) supplemented with 1 mM activated Na3VO4, one protease inhibitor mini-tablet (Thermo Scientific) and Benzonase nuclease (Millipore). For VLP production in HEK293T cells, cells were seeded in T175 flasks (4-5 flasks) at a 60% confluency, and each flask transfected with 35 μg total plasmid DNA plus 105 μl of PEI. For the viral system, the cotransfection of the MLV *gag* gene together with the receptor (Strep-tagII, HA or FLAG-tagged) plus the transporter (NA-SA/scfvHA or scfvFLAG) was made with a weight ratio of 1.0:0.4:0.6 (GAG: Transporter: Gene of interest). At 72h after transfection conditioned media was collected and adherent cells were lysed with 2 ml of RIPA buffer to be used as an expression control. For transfections in the Expi293F cell line (ThermoFisher), 100 ml of cells at 3 X10^6^ cells/ml were transfected with 100 µg of total DNA using the ThermoFisher Expifectamine transfection kit (cat #A14524) according to manufacturer’s instructions. Briefly the DNA was mixed with Opti-Plex Complexation Buffer and the expifectamine reagent and added to cells. 20 h after transfection enhancers 1 and 2 were added, and the conditioned medium collected 4 days after transfection (counting the day of transfection as day 0). For cell-based activation assays, 18 h after transfection cells were washed three times with 2 ml Ham’s F12 supplemented with 1 mg/ml BSA and serum starved in this medium for 3 h at 37°C. Each specific ligand was added in designated wells, 100 ng/ml EGF (purified as described in (65), for 5 min; and 200 nM (1.14 µg/ml) insulin (Novus Biologicals, Cat #NBP1-99193) for 30 min. Cells were washed with ice-cold phosphate-buffered saline and lysed for 30 min at 4°C in 250 µl of RIPA buffer.

### Purification of VLP/EVs

Conditioned media was clarified by two centrifugations at 4000 g for 30 min followed by two filtrations using a 0.45 µm filter unit. The media was then ultracentrifuged on a 20% sucrose cushion for 2 h at 100,000 rpm using a Tni51 rotor.

The pellets containing now the vesicles were resuspended in 50-2000 µl of TNE buffer (50 mM Tris-HCl pH 8, 100 mM NaCl, 0.1 mM EDTA) over night, except for Insulin receptor containing vesicles that the buffer was modified to 50 mM Tris-HCl pH 7, 300 mM NaCl, 0.2 mM EDTA to avoid vesicle aggregation. Absorbance at 280 nm was used to estimate if the concentration of vesicles was enough to be used for cryoEM with an OD280 of 2 being the minimal acceptable value.

The vesicles were purified by affinity purification, using the Strep-tag II fused to the N-terminus of the receptors expressed on the vesicles membranes to bind to a StreptactinXT-4flow high-capacity resin (IBA cat#2-5030-025) column. We followed the manufacturer’s protocol using 2 ml of resin per 100 ml of conditioned media. The sample was incubated in batch mode overnight at 4°C followed by rocking overnight in elution buffer at 4°C. Elution fractions of 1 ml were collected and analyzed by Western blot using the following antibodies: rabbit anti-HA Polyclonal (Thermo Fisher Scientific, Cat #PA1-985); rabbit anti-DYKDDDDK (FLAG) Tag Polyclonal Antibody (Thermo Fisher Scientific, Cat #PA1-984B); mouse anti-NWSHPQFEK Tag (Strep-tag II) Monoclonal Antibody (5A9F9); rabbit anti-EGFR (D38B1, Cell Signaling, Cat #4267), rabbit anti Insulin Receptor β-chain (Millipore Sigma, Cat #07-724); rabbit Monoclonal Antibody anti-Smoothened (E6Z5T, Cell Signaling, Cat #92981)). Rabbit anti phospho-EGFR pTyr1068 antibody (Thermo Fisher Scientific, Cat #44-788G); mouse anti-phosphotyrosine clone 4G10 (EMD Millipore, Cat #05-321); rabbit anti phospho-IR β Tyr1150/1151 (Cell Signaling Technology, Cat #2969). Rabbit anti-β-Actin (Cell Signaling Technology, Cat #4968) and rabbit anti- Hsp90α (Cell Signaling Technology, Cat #4877) were used to calibrate protein loading concentrations. Goat anti-rabbit- 680RD (Li-Cor, Cat #926-68071) and Goat anti-mouse IgG2b-800CW (Li-Cor, Cat #926-32352) were used as secondary antibodies and detected using a Li-Cor Odyssey Clx Near IR imaging system.

Positive affinity-column elution fractions were combined and ultracentrifuged through a 20% sucrose cushion in PBS for 2h at 100,000 rpm using a Beckman SW 32 Ti rotor. Pellets were resuspended in 50-200 µl of TNE overnight with gentle agitation at 4°C. The concentrated vesicles were quantified by spectrophotometry by optical density at 280 nm and screened by negative stain electron microscopy using Formvar carbon supported copper grids, 300 mesh (Electron Microscopy Sciences #FCF300-Cu-50 241141). Four μl of sample at ∼OD280 ≥ 2.0 was applied to the grid. After 2 min, the sample was blotted and immediately washed twice in water for 30 s before incubation with 4 µl of 2% Uranyl Acetate at pH 4.5 for 45 s. The grid was then blotted and air-dried. Images were collected using an FEI JEOL NEOARM scanning transmission electron microscope operated at 200 kV equipped with an EDS detector. The vesicles were then aliquoted, and flash frozen with liquid nitrogen, stored at -80 °C. In vitro phosphorylation of the EVs was tested by incubating 10 μl of purified EVs with 1 mM ATP, 1mM Na3VO4, 20 nm HEPES, 50 mM MgCl2 with or without of 1 μg/ml EGF incubated at room temperature for 5 minutes. Loading buffer was added and samples analyzed by Western Blot.

### Mass spectrometry sample preparation

VLPs or EVs in TNE buffer (50 mM Tris-Cl pH 8.0, 200 mM NaCl, 0.1 mM EDTA) were prepared for mass spectrometry using the membrane protein-optimized method of Lin et al. (66). Briefly, an equal volume of 4% SDS was added (for 2% final concentration), samples were boiled 5 minutes and proteins precipitated with 6 volumes ice cold acetone for at least 4 hours 4°C. The precipitate was pelleted at 16,000 x g for 15 minutes at 4°C, the supernatant was discarded, and the pellet washed twice with 400 µl cold acetone and re-centrifuged as before. Care was taken to remove all but ∼20 µl of supernatant in each cycle to preserve the delicate pellet. The washed pellet was air dried and resuspended in 100 µl 1% sodium deoxycholate, 50 mM ammonium bicarbonate with bath sonication (2 x 10 minutes). Proteins were reduced (TCEP to 5 mM, 56°C, 45 minutes) and alkylated (iodoacetamide 25 mM, 45 min room temperature in the dark). Alkylation was quenched with 12 mM DTT and proteins digested overnight at 37°C with 0.5-1 µg MS-grade trypsin (Pierce #90058). Trypsin was inactivated and sodium deoxycholate precipitated by addition of formic acid to 1% followed by centrifugation at 16,000 x g at 4°C for 10 minutes. The supernatant was transferred to a fresh tube and the volume adjusted to 250 µl prior to ultrafiltration in in a pre-washed Vivaspin500 (Sartorius) or AmiconUltra (Millipore) 10 kDa filtration unit. Filtrate was dried and resuspended in 20 µl 0.1% TFA for desalting using a ZipTip (Millipore #ZTC18S096), eluted in 10 µl 50% acetonitrile and 0.1% TFA, dried, and resuspended in 18-20 µl 5% acetonitrile, 0.1% formic acid for LC/MS. Two to three 5 µl injections of each sample were performed on a minimum of two replicate samples for each protein of interest.

### Mass spectrometry data collection and analysis

Mass spectra were collected using a Thermo Orbitrap Fusion Lumos Tribrid mass spectrometer and a data-dependent 75- minute top speed collection method with a 60 minute 3-40% acetonitrile gradient, dynamic exclusion after one observation for 15 seconds, and stepped HCD (27/30/33). RAW files were individually processed in Proteome Discoverer 2.5 using the PWF Tribrid_Basic_SequestHT workflow with the Percolator node for PSM validation and the following settings: trypsin digestion with up to 2 missed cleavages, static carbamidomethyl modification of cysteine, and dynamic modifications of oxidized methionine, and protein N-terminal modifications of Met-loss, Acetyl, or Met-loss + Acetyl. For spectral searches, FASTA files containing sequences of the synthetic tagged transporter and protein of interest were included along with the standard Uniprot human proteome and common contaminants. Data including FASTA files are available from the MassIVE data repository # . Initially, each RAW file was processed to produce an msf file, and the msf files from samples to be compared were reprocessed as needed using the CWF_Basic workflow with Merge Mode “Do Not Merge” to create a single Consensus file for comparative analysis. Results were filtered to remove contaminants and proteins identified by a single PSM. For comparing relative protein abundances in control VLP/EVs vs VLP/EVs containing the protein of interest, differential protein enrichment was computed using the RUVr edgeR-quasi-likelihood model implemented in Degust (67), requiring a minimum count of 2 PSMs in at least 2 samples, and normalizing by Trimmed Mean of M-values. Absolute abundances of membrane proteins were estimated using the label-free iBAQ analysis implemented in MaxQuant, identifying transmembrane proteins based on Uniprot annotations, and computing histograms with Perseus.

### Cryo-ET sample preparation

Concentrated EVs were mixed with 10-nm colloidal gold (in PBS solution) in a 10:1 ratio. 3 µl of the solution was then added to a glow-discharged ultrafoil gold grid (Quantifoil 2/2, 200 mesh, Ted Pella #687-200-AU-50).

Grids were plunge frozen into liquid ethane by double-side blotting using a Vitrobot cryo plunger (Thermo Fisher) and stored in liquid nitrogen until imaging.

### Cryo-ET data collection

Cryo-ET data collection was performed essentially as described previously (33). Cryo-grids containing the EVs were loaded into an FEI Titan Krios transmission electron microscope operated at 300 kV, and images were recorded on a Gatan K3 Biocontinuum Imaging Filter (Gatan) direct detection camera in counting mode with a 20 eV energy slit in zero-loss mode. Tomographic tilt series between −60° and +60° were collected with serialEM (68) using a dose-symmetric scheme (69) with a 3° angular increment. A total dose of 120 e^-^/A^2^ per tilt series was distributed evenly among 41 tilt images. The nominal magnification was 53,000 X, giving a pixel size of 1.69 Å on the specimen. The defocus range was between -4 µm and -5 µm and 8 frames were saved for each tilt angle. All data acquisition parameters are listed in Supplemental Table 1.

### Cryo-ET data processing

The preprocessing of the tilt series was done in RELION5 (70, 71) using *relion --tomo*. The movies and mdoc files were imported, the frames were motion-corrected using MotionCor2 (72) and the contrast transfer function (CTF) was measured using the CTFFIND4 package (73). After poor-quality tilt images were excluded, the tilt series stacks were aligned automatically with the 10-nm fiducials, an IMOD (74) function adapted within RELION5. The 10-nm gold fiducials were removed using *ccderaser* and *newstack* functions from the motion-corrected micrographs. The final tomogram reconstruction was carried out using the gold fiducial-based alignment and the fiducial-erased micrographs. For better visualization, tomograms were low-pass filtered to 50 Å using e2proc3d function from EMAN2 (75) and tomographic slices were visualized with IMOD (74).

### Extraction of EGFR dimers from tomograms

The initial steps of subtomogram alignment and averaging were done with the subTOM package (Dustin Morado subTOM: https://github.com/DustinMorado/subTOM), which was originally implemented using MATLAB (MathWorks) scripts derived from the TOM (76) and AV3 (77) packages as described previously (33).

To generate an initial template model of the EGFR protein from the EV surface, 160 EGFR particles were manually picked from 7 EVs that were down-scaled by 4x binning of the voxels (pixel size of 6.76 Å). The 160 EGFR particles’ initial Euler angles (2 out of 3) were determined based on the vector between two points, one on the head of the EGFR and one on the membrane where the EGFR anchors, respectively. The 160 EGFR were iteratively aligned to one another for three iterations, revealing a two-fold symmetrical EGFR dimer. Hereafter, this template and two-fold symmetry was used during the subsequent alignment and averaging process.

We tried to automatically locate the rest of the EGFR dimers on all the EVs with the existing initial template model by applying a previous oversampled spherical grid of points approach that worked for SARS-CoV-2 spike protein (33). However, due to the much-lower molecular weight (150 kDa vs. 500 kDa), lower-symmetry (C2 vs. C3), and the crowdedness of EGFR proteins on the EV surface, automatic template matching the EGFR dimers on EVs failed.

We then manually picked 1602 EGFR dimers from 22 tomograms using the two-point approach, used earlier to determine the initial template model. Subtomograms were aligned against the low-resolution template (from the above average of 1602 EGFR dimers). Visual inspection of the tomograms using the Place Object Chimera Plugin (78) confirmed that subtomograms selected in this manner corresponded to EGFR dimers on the EV surface. Iterative alignment and averaging were performed at a binning of 4 (pixel size of 6.76 Å) in subTOM and resulted in a 32 Å averaged dimer. Subtomograms were divided into two halves based on the tomogram number. From this point on the two halves were processed independently in RELION5 (71).

### Subtomogram averaging of EGFR in RELION

Subsequent processing was performed in RELION5 (70, 71) as described in (33). For this purpose, subtomograms were reconstructed from the gold fiducial-removed micrographs after motion correction. Using dedicated python scripts, the EGFR positions in the 3D tomograms from subTOM were converted into 2D positions and defocus values in the corresponding tilt series images, as well as Euler angles in the RELION starfile convention. Individual subtomograms were reconstructed at a 3x down-scaled pixel size of 5.07 Å in a cropped box size of 96 voxels.

Standard 3D auto-refinement was performed with C2 symmetry and a soft-edged mask around the EGFR dimers, using a 60 Å low-pass filtered map generated from subTOM alignment parameters as initial reference. 3D classification was applied to sort the particle heterogeneity, and 843 of the 1,602 particles were kept for further refinement. Finally, a 15 Å consensus map was calculated for the EGFR dimers.

*Subtomogram averaging of ICR.* The averaging process for the ICR is similar to the EGFR ectodomain, with a two-point approach mentioned above. Because the crowded ICR particles are generally too noisy to be resolved and averaged, we chose to pick only isolated and well-defined ICR particles for averaging, and an average was generated from 16 particles. Due to the low number of particles and small protein size, we did not proceed with iterative alignment.

*Quantification of EV diameter.* For EV diameter measurements, EV diameters were quantified from cryo-ET tomographic sections that were 4x binned and low-pass filtered to 50 Å tomograms (EMAN2 *e2proc3d.py*) (75) for better visualization. Quantification of EV diameter was done using IMOD by adding open contour points on each complete EV (n = 167) and the diameters were calculated. Note, incomplete EVs were not considered for quantification. The *imodinfo* utility was used to extract the virion diameter information from the saved model files.

*Segmentation of EV and EGFR.* Density map segmentation and surface rendering were conducted using the Volume Tracer and Color Zone tools in UCSF ChimeraX (79). Markers were first added in ChimeraX to designate the ECR, membrane, and ICR regions within the tomograms. These regions were then segmented into different maps using the Color Zone tool, with each region assigned a unique color for clear visualization.

*Radial average density profile.* The “Radial Profile Angle” plugin in ImageJ (80) was employed to calculate the radial average density profile of regions with and without ICR and ECR in EVs. The plugin was used to identify the center, define the starting angle, and set an integration angle of 25 degrees for each region. The radial average density was then calculated, and the resulting data were exported to MATLAB for further analysis.

### Data Sharing Plan

The EGFR ectodomain structure will be deposited at Electron Microscopy Data Bank (EMDB) prior to publication. The sequences encoding all transporter proteins will be deposited in GenBank prior to publication. All plasmids will be made available upon request or deposited in Addgene. Mass spectrometry data including FASTA files have been deposited in the MassIVE data repository (MSV000096518).

## Acknowledgements

We thank Evan Schwartz from the Sauer Structural Biology Lab for technical assistance with cryo-EM data collection. We thank Xun Zhan of the Texas Materials Institute EM facility and Michelle Mikesh of The University of Texas at Austin Center for Biomedical Research Support for technical assistance with negative-stain EM. We thank Nancy Carrasco for supplying the NIS clone. Funding for these studies was provided by awards from the Cancer Prevention and Research Institute of Texas (RR160023) and the Wojcicki Foundation (#LC029) (DJL); Cancer Prevention and Research Institute of Texas (RR230050) (ZK); and the National Institute of General Medical Sciences (R35GM122480), Army Research Office (W911NF-12-1-0390), and Welch Foundation (F-1515) (EMM). The Sauer Structural Biology Laboratory is supported by the University of Texas College of Natural Sciences and by award RR160023 from the Cancer Prevention and Research Institute of Texas.

## Disclosure and competing interest statement

The authors declare no competing interests.

---

**Figure S1.**
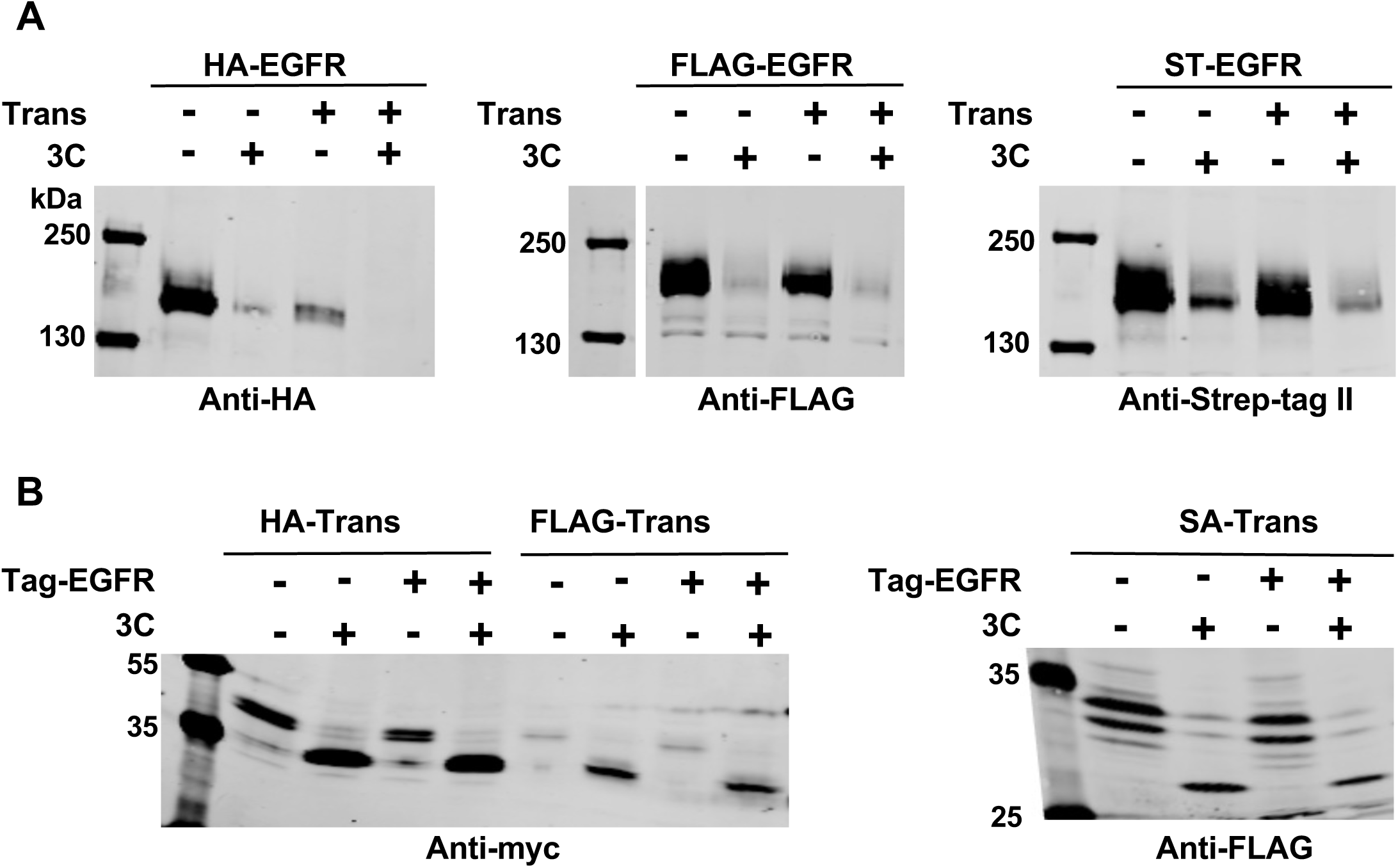
Extracellular regions of transporter proteins (Trans) and epitope tags can be removed by Rhinovirus 3C protease. (A) Western blots of cell lysates of HEK 293T cells transfected with genes encoding EGFR labeled at its N-terminus with a hemagglutinin peptide (HA-EGFR), a FLAG peptide (FLAG-EGFR), or the Strep-tag II peptide (ST-EGFR) either with or without co-transfection with the cognate transporter. (B)The top row shows cleavage of the tag from the EGFR N-terminus showing efficient removal of epitope tags by 3C. (B) Western blots of HEK 293T cells transfected with transporter proteins with anti-HA (HA-Trans), anti-FLAG (FLAG-Trans), or Strep-tactin (SA-Trans) modules showing efficient cleavage of the myc-labeled (HA-Trans and FLAG-Trans) or FLAG-labeled (SA-Trans) transporters by 3C protease. The probing antibody is indicated below each blot.

**Figure S2.**
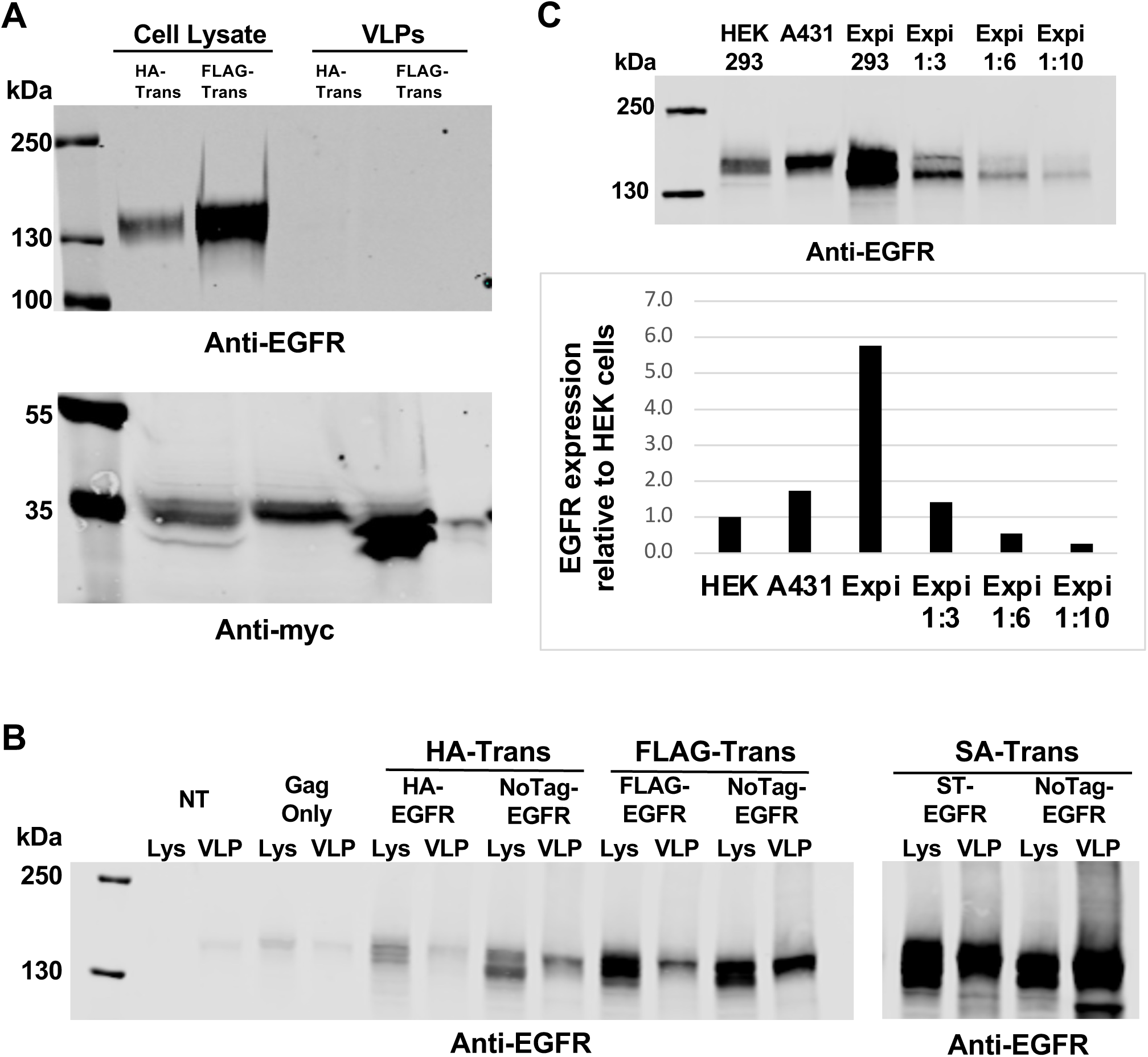
The transporter protein is needed to transport tagged proteins to VLPs derived from HEK 293T cells but not Expi293 cells. (A) Western blots of cell lysates and VLPs derived from HEK293 cells transfected with untagged EGFR, Gag, and either the HA (HA- Trans) or FLAG transporter (FLAG-Trans). The probing antibody is indicated below each blot. EGFR is not present in VLPs when untagged. (B) Western blots of cell lysates and VLPs derived from Expi293 cells transfected with tagged and untagged versions of EGFR, Gag, and cognate transporters. Untagged EGFR is transported to VLPs when expressed in Expi293 cells. (C). Western blots of lysates from A431 cells and HEK 293T and Expi293 cells transfected with the EGFR gene. The amount of lysate loaded was normalized to the protein concentration using a BCA assay.

**Figure S3.**
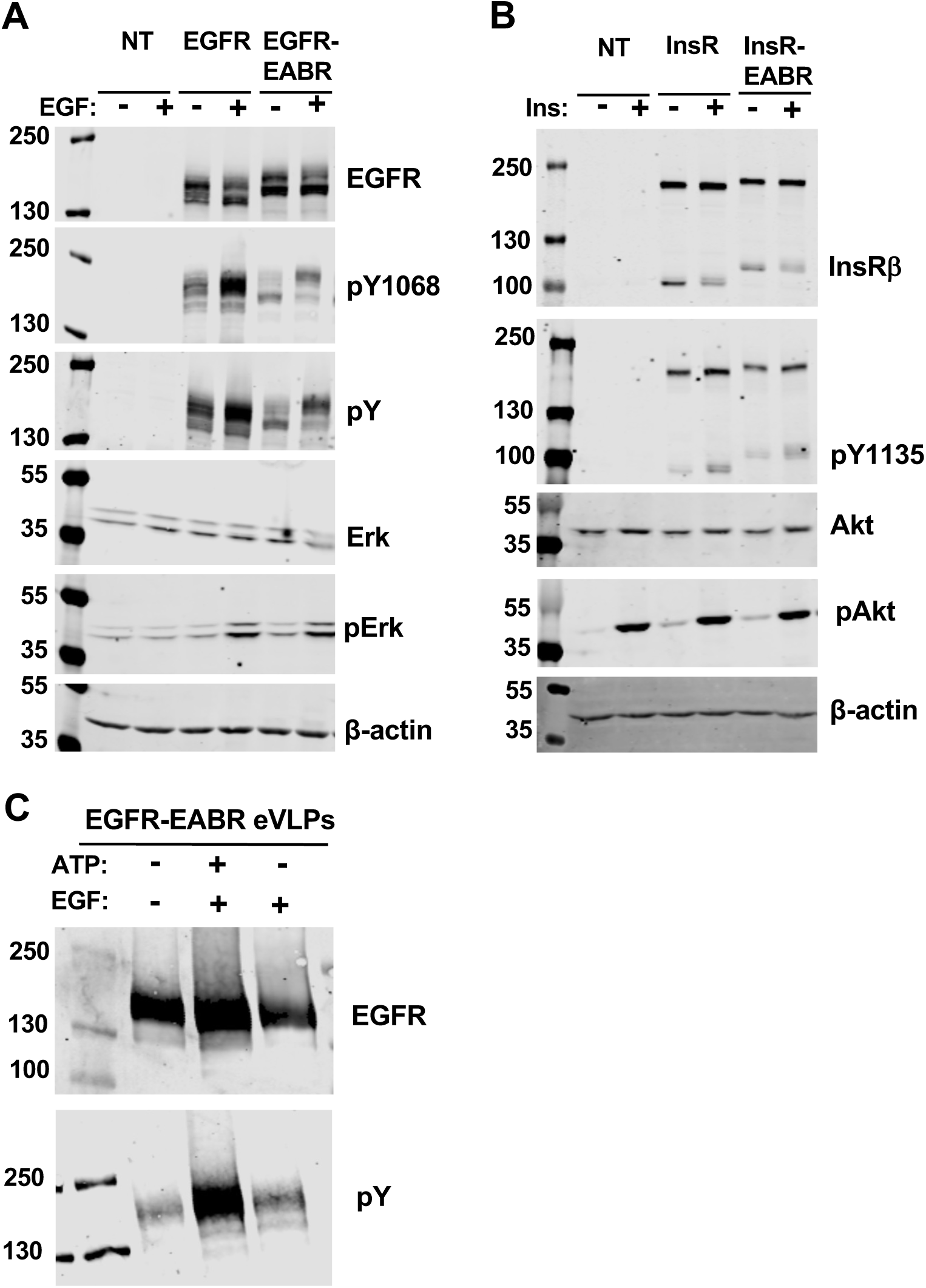
Addition of EABR sequences to EGFR and InsR does not inhibit activation in cells or EGFR in EVs. (A) Western blot analyses of lysates of CHO cells either not transfected (NT), transfected with a gene encoding EGFR (EGFR), or transfected with a gene encoding EGFR with a C-terminal EABR sequence (EGFR-EABR) and either treated (+) or not treated (-) with EGF. (B) Western blot analyses of lysates of CHO cells either not transfected (NT), transfected with a gene encoding Insulin Receptor (InsR), or transfected with a gene encoding InsR with a C-terminal EABR sequence (InsR-EABR) and either treated (+) or not treated (-) with Insulin. (C) Western blots of EGFR-EABR containing eVLPs showing increased receptor phosphorylation following addition of ATP and ligand. The probing antibody is indicated beside each blot.

**Figure S4.**
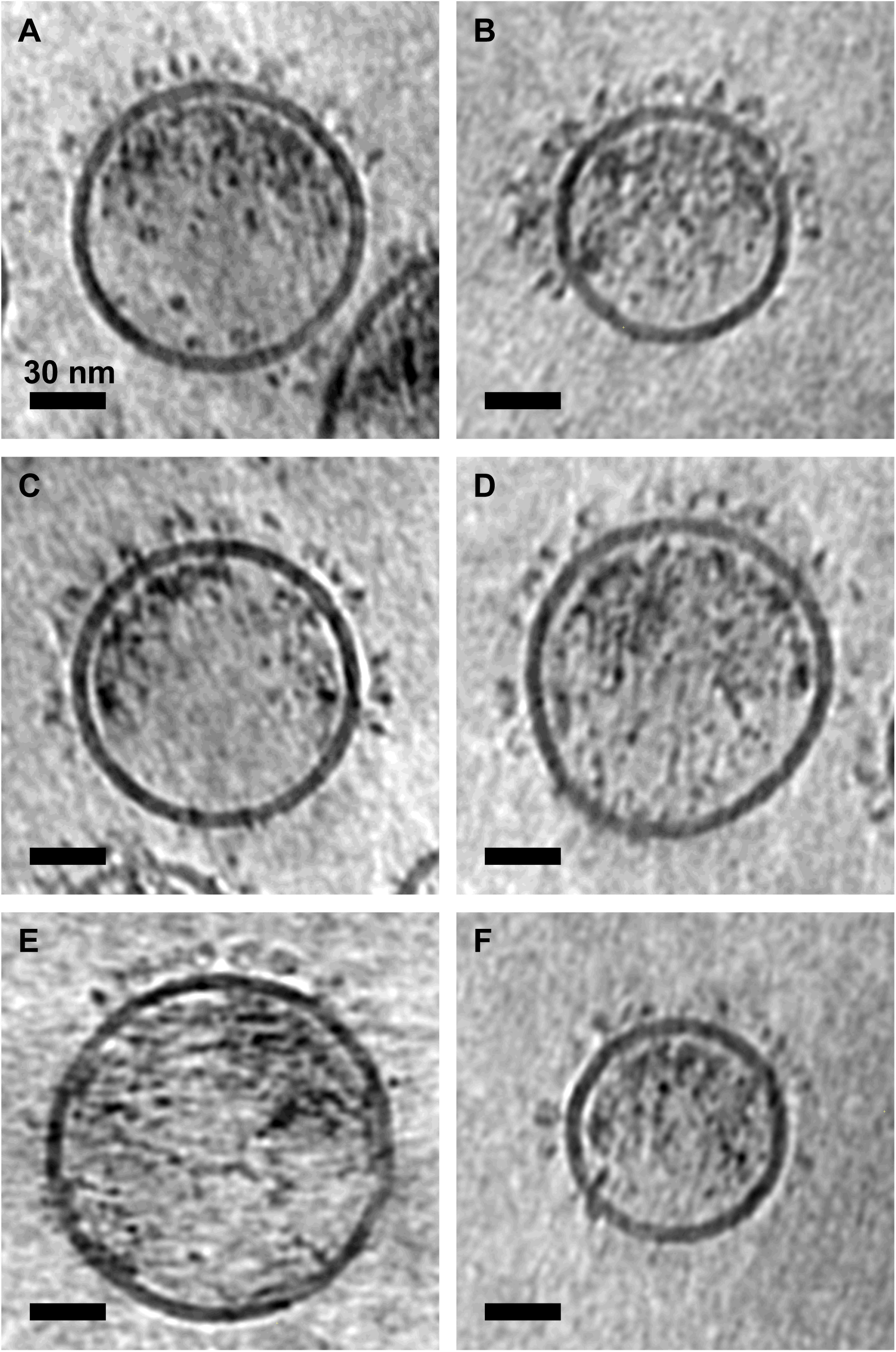
Sections of tomograms of EGFR-containing EVs are shown. The tomogram in panel A is the same tomogram shown in Figure 3A and is reproduced here for comparison. All scale bars are 30 nm.

**Figure S5.**
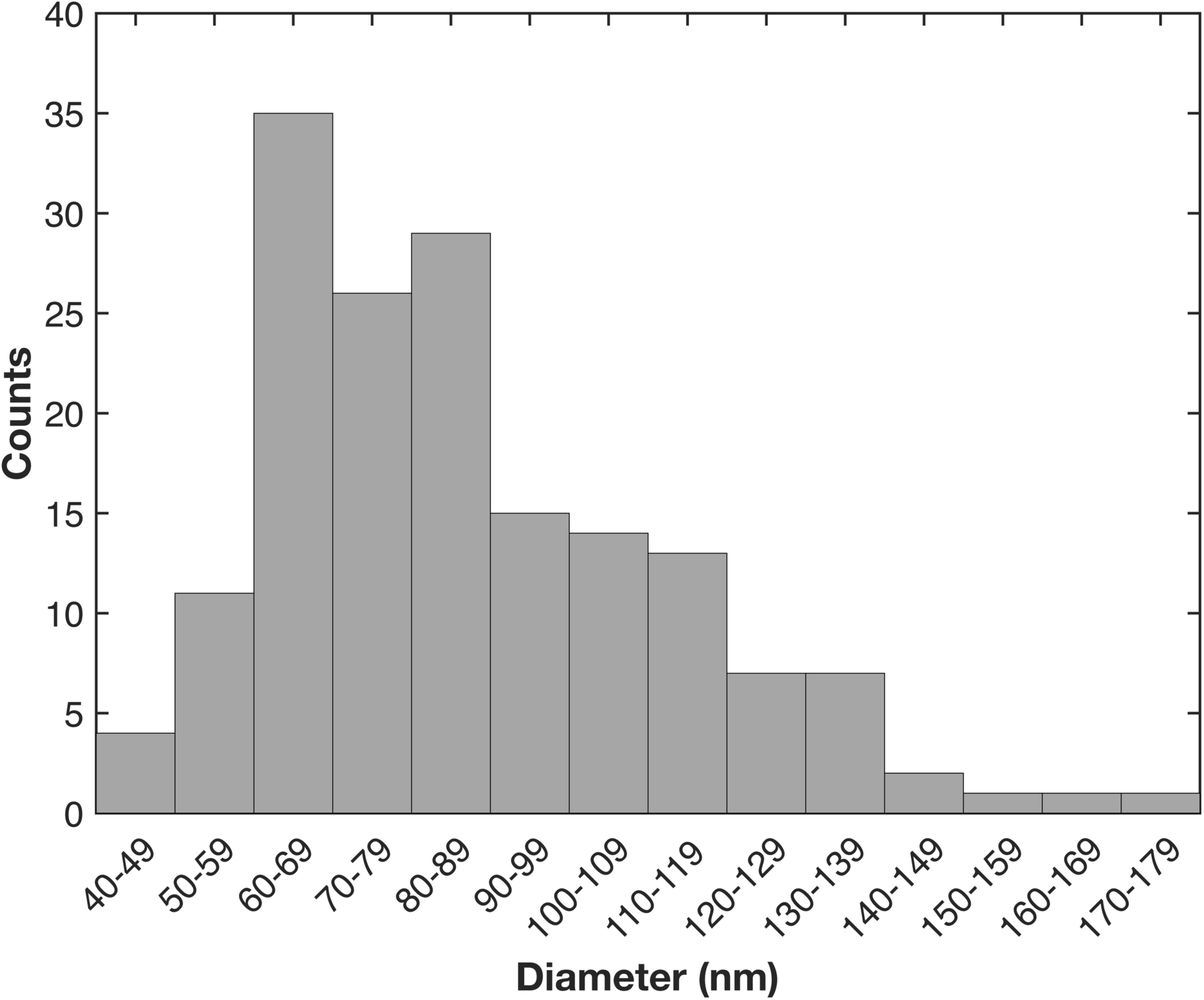
Histogram of diameter of EGFR-containing EVs in tomograms. A total of 167 EVs were measured.

**Figure S6.**
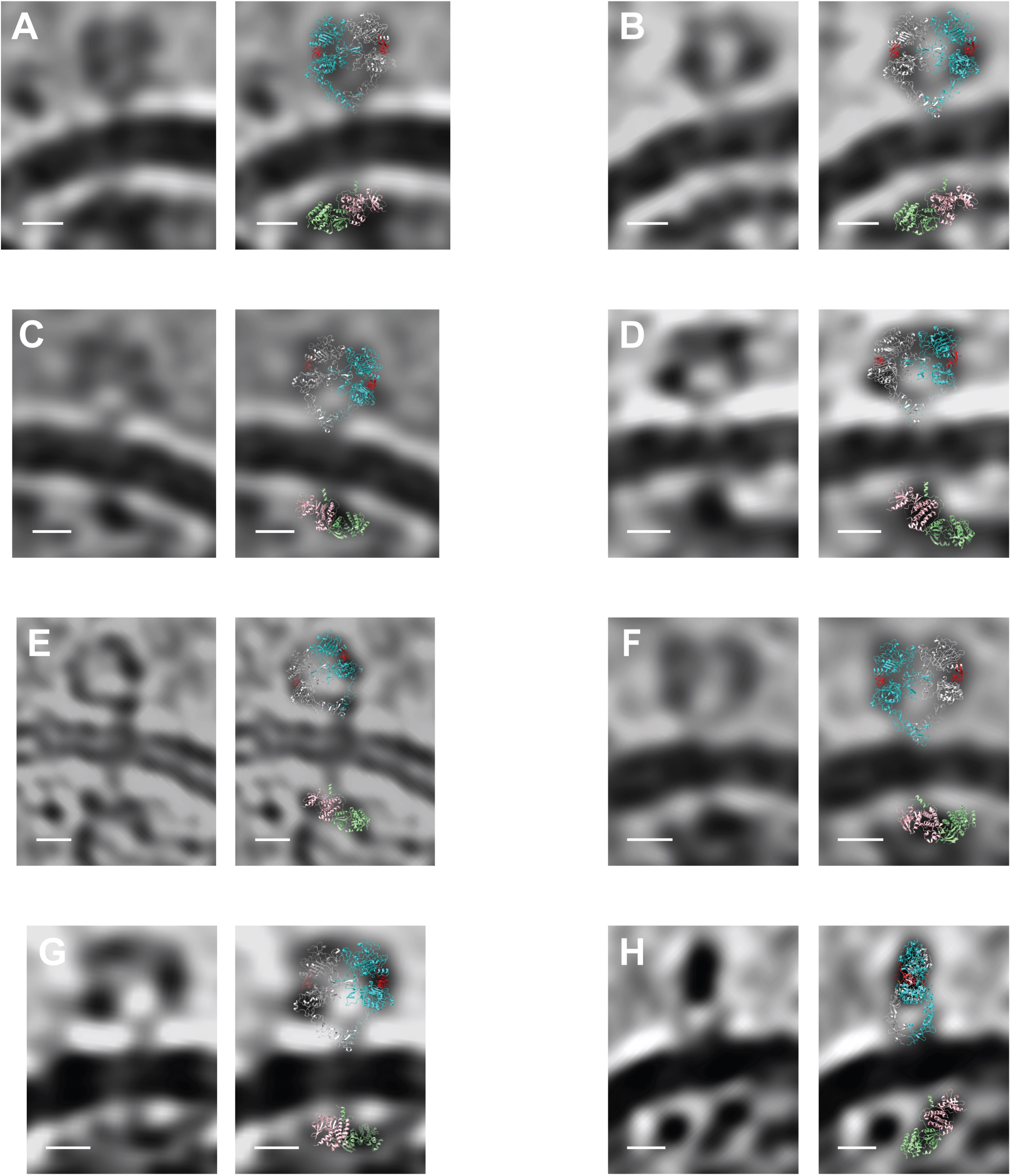
Images of individual EGFR molecules from tomograms of EGFR-containing EVs are shown alongside the same image with the crystal structures of the EGFR extracellular region (RCSB 3NJP; one EGFR subunit colored cyan and the other light gray with EGF colored red) and the asymmetric EGFR kinase dimer (RCSB 3GOP; one kinase subunit is colored light pink and the other light green). All scale bars are 5 nm.

**Figure S7.**
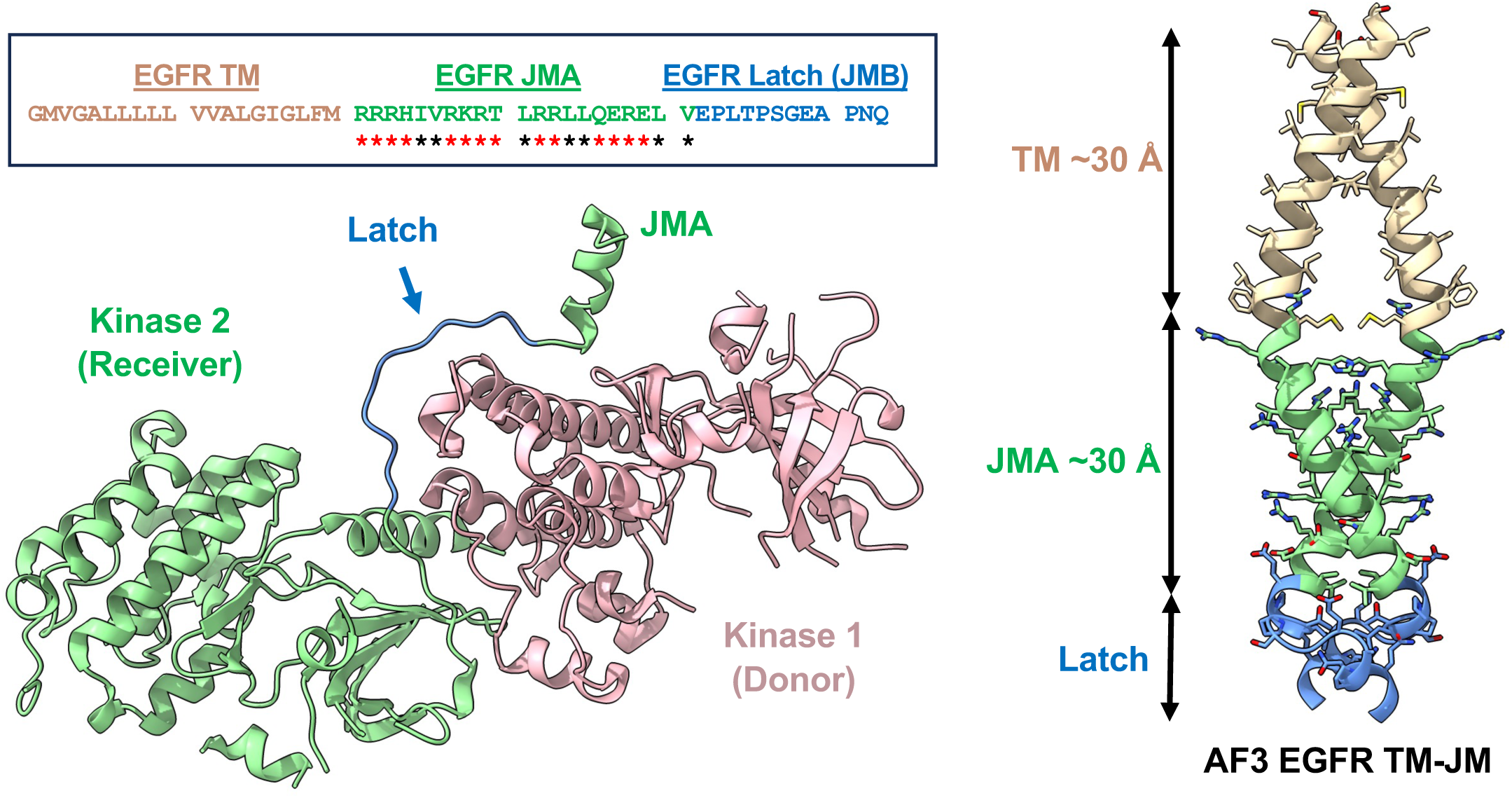
The juxtamembrane A region of EGFR is predicted by Alphafold 3 to form a coiled-coil like structure. The sequence of the EGFR transmembrane (TM) and juxtamembrane (JM) regions is shown in the inset with red and black asterisks indicating hydrophilic and hydrophobic residues in the JMA region, respectively. A ribbon diagram of the asymmetric dimer of the EGFR kinase (RCSB 3GOP) is shown (left) with the donor kinase colored light pink and the receiver light green. The JM latch and JMA regions are colored light blue and light green, respectively. An Alphafold 3 model of a dimer the EGFR TM and JM regions is shown (right) with the TM region colored wheat, the JMA region colored light green, and the Latch (JMB) region colored light blue. The C-terminal 3 turns of the JMA region of the Alphafold3 model match the 3 turns of the JMA helix helix visible in the kinase crystal structure.

**Figure S8.**
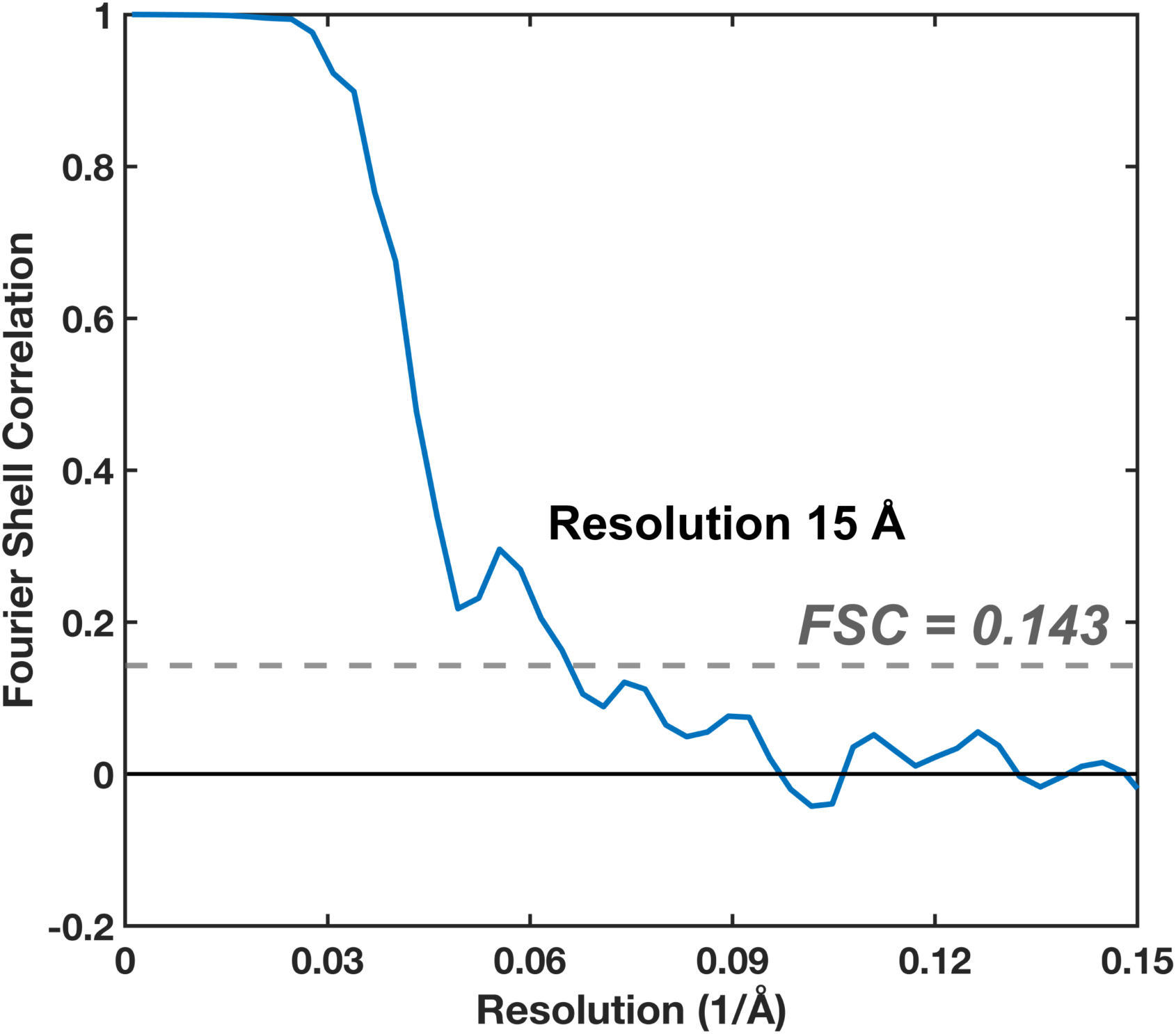
Fourier shell correlation plot for the subtomogram average of 843 EGFR extracellular region particles.

**Supplementary Table 1.**
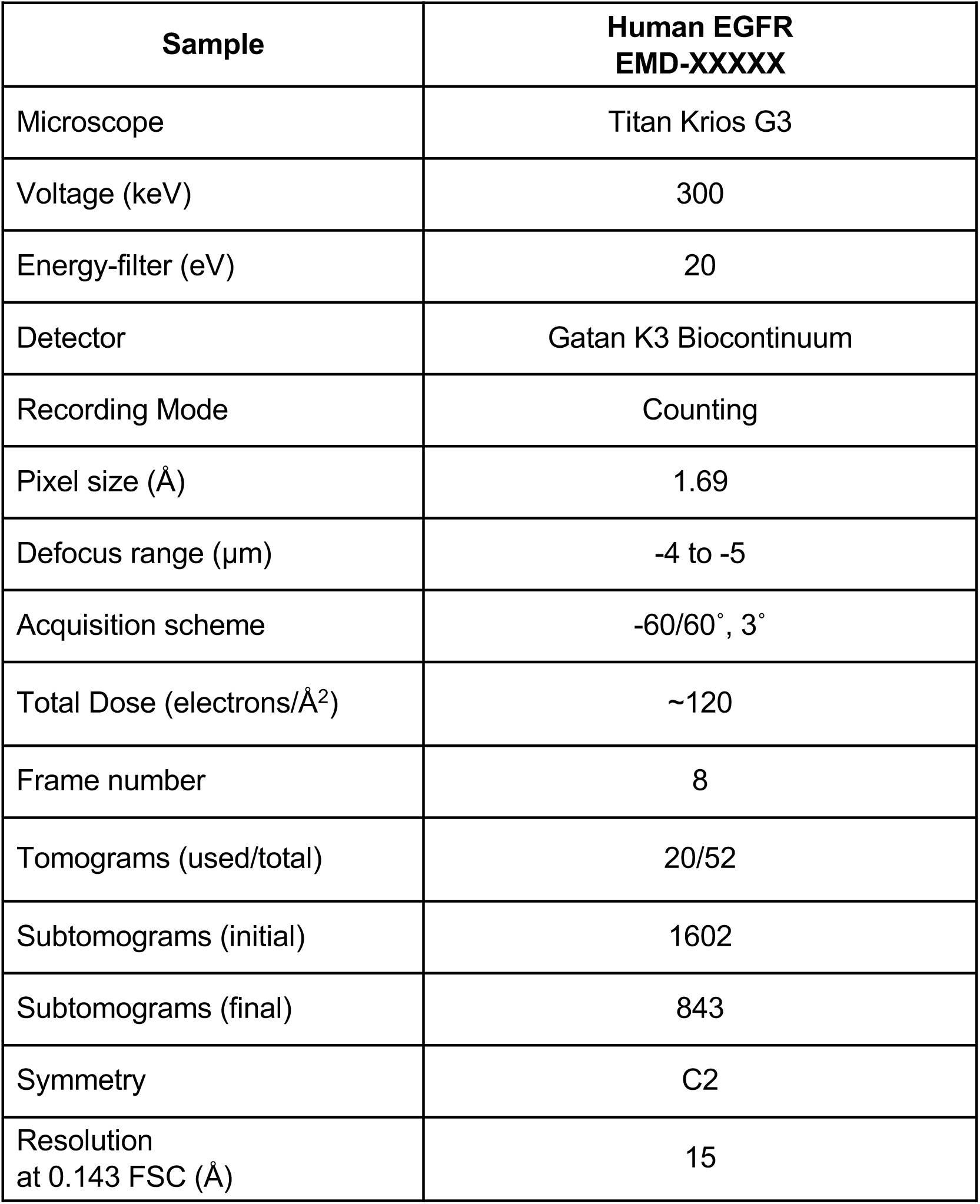
Cryo-ET data acquisition and image processing.

| **Sample** | **Human EGFR**<br>**EMD-XXXXX** |
| --- | --- |
| Microscope | Titan Krios G3 |
| Voltage (keV) | 300 |
| Energy-filter (eV) | 20 |
| Detector | Gatan K3 Biocontinuum |
| Recording Mode | Counting |
| Pixel size (Å) | 1.69 |
| Defocus range (µm) | -4 to -5 |
| Acquisition scheme | -60/60°, 3° |
| Total Dose (electrons/Å<sup>2</sup>) | ~120 |
| Frame number | 8 |
| Tomograms (used/total) | 20/52 |
| Subtomograms (initial) | 1602 |
| Subtomograms (final) | 843 |
| Symmetry | C2 |
| Resolution<br>at 0.143 FSC (Å) | 15 |

## Notes

### Competing Interest Statement

The authors have declared no competing interest.

